# Adaptive immune responses to vaccination reflect social status at first exposure in female rhesus macaques

**DOI:** 10.64898/2026.03.03.709117

**Authors:** Joao Barroso-Batista, C. Ryan Campbell, Noah D. Simons, Paul L. Maurizio, Raúl Aguirre-Gamboa, Anne Dumaine, Mari Shiratori, Tawni Mondragon, Cary Brandolino, Vasiliki Michopoulos, Jenny Tung, Luis B. Barreiro

## Abstract

Social gradients are consistently associated with variation in health outcomes, including infectious disease. However, distinguishing between social gradients in antigen exposure versus social gradients in susceptibility remains challenging. Here, we use a nonhuman primate model for chronic social stress to investigate how social status affects the influenza vaccine-induced adaptive immune response. We first manipulated the social status of female rhesus macaques and then tested the response to influenza antigens in naïve individuals and after secondary exposure. Higher social status at the time of first exposure, but not at the time of secondary exposure, predicted stronger antibody responses to both exposures. Social status also drove gene expression differences in adaptive immune pathways, and genes that predicted the magnitude of the antibody response overlapped with those linked to social status. Thus, social gradients shape the adaptive immune response in a temporally dependent manner, with particular sensitivity at the time of initial antigen exposure.

## Introduction

Social factors, including socio-economic status (SES), social integration, and early-life adversity, are among the strongest and most consistent predictors of health described for human populations to date (*1–7*). Social gradients affect all ten leading causes of death in the United States (*8*), including cardiovascular disease (*9*), cancer (*10*), and diabetes (*11*). These observations have led some researchers to suggest an underlying, partially shared etiology: from social disadvantage to chronic social stress, from chronic stress to immune dysregulation, and from immune dysregulation – particularly chronic inflammation – to elevated risk of multiple disease types (*12–18*). In support of this idea, various measures of social disadvantage in humans predict elevated markers of inflammation (*19–21*), altered immune cell composition (*22*), and impaired immune function (*23*). For example, multiple large cohort studies report an association between low SES (measured by education, occupation, or income) and increased levels of circulating inflammatory markers, such as CRP, IL-6, TNF-a or fibrinogen (NHANES IV (*24*), Framingham Offspring (*25*), CARDIA (*26*), Whitehall II (*27*, *28*)).

Research on social gradients in health has a strong focus on non-communicable diseases, particularly cardiovascular disease (*9*, *29–32*). However, if these gradients reflect underlying differences in immune function, they should also shape susceptibility to infectious disease. Consistent with this idea, socially disadvantaged individuals experience a higher burden of multiple infectious diseases. HIV diagnosis rates are approximately four-fold higher in high-poverty census tracts (*33*), influenza-related hospitalization rates are about two-fold higher in high-poverty counties (*34*) and, during the COVID-19 pandemic, COVID-19-related mortality increased by roughly 14 percent for each 0.1-point increase in the Social Vulnerability Index (*35*). Socially disadvantaged individuals also acquire latent viral infections (e.g., cytomegalovirus, Epstein-Barr virus, and herpes simplex virus) earlier and exhibit poorer control of these infections throughout life (*36–39*). Accordingly, socioeconomic measures such as education and income are among the strongest predictors of pathogen burden across a diverse set of infectious agents (*40–43*).

Although social gradients in infectious disease are strongly shaped by differences in exposure and health care access (*44–48*), two lines of evidence suggest that they might also be affected by systematic differences in underlying immune physiology. First, chronic caregiving-related stress predicts weakened antibody responses to influenza and pneumococcal vaccines relative to sex-, age- and socioeconomically matched controls (*49–54*). This difference has been suggested to emerge from the detrimental effect of long-term stressors on the development of a typical antibody response (*55*). Second, experimental inoculation with standardized titers of respiratory viruses, including coronavirus, respiratory syncytial virus, influenza, and rhinovirus, reveals that multiple forms of social disadvantage predict conversion to clinical symptoms (*56*). Specifically, socially isolated individuals, those experiencing chronic stress at the time of inoculation, and those who had experienced high levels of past childhood adversity fared more poorly (*57–62*). Together, these results are consistent with the idea that social disadvantage not only alters the likelihood that individuals will be exposed to infectious pathogens, but that the immune response *once infected* also varies as a function of social experience.

However, even if pathogen exposure can be experimentally controlled in studies of humans, exposure to meaningful differences in social adversity and advantage cannot. Individuals who differ in socioeconomic status, social integration, or other dimensions that shape social gradients also differ in their history of past pathogen exposure, vaccination uptake, and lifestyle (*44*, *46–48*, *63*, *64*). Animal models for the social determinants of health can partially overcome this limitation. Indeed, consistent with the idea that chronic social stress is causal to differences in infectious disease outcomes, social stress alters the outcome of experimentally controlled viral infections in lab mice, and mice who experience social disruption exhibit consistent changes in physiology, immune cell composition, and cytokine and antibody responses relative to non-stressed controls (*65–69*). In nonhuman primates, where the physiology of the immune system more closely recapitulates that in humans, social status, social isolation, and social support have all been suggested to shape the immune response to viral infections (*70–76*). For example, in both long-tailed and rhesus macaques, low social status corresponds to a higher likelihood of infection, higher viral loads, and/or shortened post-infection lifespan following controlled exposure to adenovirus or simian immunodeficiency virus (*58*, *71–73*), whereas a higher rate of affiliative social behavior is associated with protection (*71*). Together, these studies extend reports of social gradients in immunity in humans to animal models observable in a controlled setting. A limitation of such studies, though, is that they often use stress paradigms that animals are unlikely to encounter in nature (e.g., repeated bouts of group instability (*77*)) or, like the Pittsburgh Cold Studies in humans (*56*), manipulate exposure to infectious agents but not the social environment itself. Thus, whether ethologically relevant, chronic stress-associated variation in the social environment is causally important to the response to infectious agents remains incompletely resolved.

Here, we address this question by drawing on experimental manipulations of dominance rank in female rhesus macaques (*78–80*). In this species-sex combination, dominance rank is a linear measure of social status that can remain stable for many years and between generations (*78*). Low status individuals tend to groom and be groomed less often (the major currency of social affiliation in this species) and are also more frequently targets of aggressive behavior from other animals (*81*). Consequently, low status females often experience elevated glucocorticoid levels and markers of glucocorticoid resistance, consistent with chronic, socially induced stress (*82*, *83*). Studies that directly manipulate rank hierarchies in newly formed social groups show that differences in social status are causal to an array of behavioral, physiological, and molecular outcomes (*82*, *84*, *85*), including differences in peripheral immune cell composition, cytokine levels, and gene expression and chromatin accessibility in immune cell types (*74*, *79*, *80*, *86–89*). These findings support links between chronic social disadvantage, chronic social stress, and changes in immune function, but have not yet shown that these changes in turn influence immunological outcomes directly relevant to health.

To do so here, we experimentally manipulated the social status of 47 adult rhesus macaque females in ten social groups, following a previously established paradigm (*78–80*, *88*). We then investigated three sets of questions. First, we asked whether systematic differences in social status *per se* influence the immune system’s ability to mount protective antibody responses to vaccination with influenza A and B antigens, an outcome with direct relevance to social gradients in morbidity and mortality in humans (*34*, *47*, *90–93*). Second, after one year, we rearranged the structure of the social groups in a way that scrambled the social status of study subjects. This approach allowed us to test the implications of social advantage and disadvantage for longer-term adaptive immune function, under the hypothesis that social status at the time of first antigen exposure should be most consequential for shaping the antibody response. This possibility might help explain why early life social disadvantage has been observed to continue predicting immune dysregulation later in life (*94*), as well as why, in previous studies of rhesus macaques, an individual’s social history continues to predict immune cell gene expression after the social environment has changed (*88*). Finally, we investigated whether the relationship between social status and vaccine responsiveness is related to social gradients in gene regulation, which have been reported both in humans and nonhuman primates (reviewed in (*95*, *96*)). These studies have often been interpreted as evidence for compromised immune function in socially disadvantaged individuals. Here, our analysis investigates the relationship between changes in immune gene regulation and the response to influenza antigens directly.

## Results

### Social status influences the response to influenza vaccination in naïve rhesus macaques

To investigate how social status-driven variation in the social environment influences the response to influenza vaccination, we first experimentally manipulated the dominance ranks of 47 adult female rhesus macaques housed at the Emory National Primate Research Center (ENPRC) in ten distinct social groups (n = 3 – 5 animals per group; Data S1). To do so, we used an established protocol in which order of introduction into a newly constructed social group is the primary predictor of subsequent dominance rank (*80*) (Fig. 1A). Co-housed females were socially unfamiliar to one another prior to introduction and none were close kin. As in previous studies, earlier introduction corresponds to higher dominance rank (Pearson’s *r* = −0.51, *p* = 1.9 x 10^-4^), which we represent here using continuous Elo ratings (*97*) (high values correspond to higher rank/social status). As expected, variation in Elo rating translates into marked differences in social experience: agonistic behavior is more frequently targeted towards lower ranking females than higher ranking females (Fig. 1B top panel, Pearson’s *r* = −0.78, *p* < 2.1 x 10^-10^), while higher ranking females engage in affiliative grooming behavior more often than lower ranking females (Fig. 1B bottom panel, Pearson’s *r* = 0.48, *p* = 6.6 x 10^-4^). Once established, dominance hierarchies remained highly stable over time, consistent with social dynamics typical for rhesus macaque females (stability index S = 0.997, see Material and Methods).

**Fig. 1.**
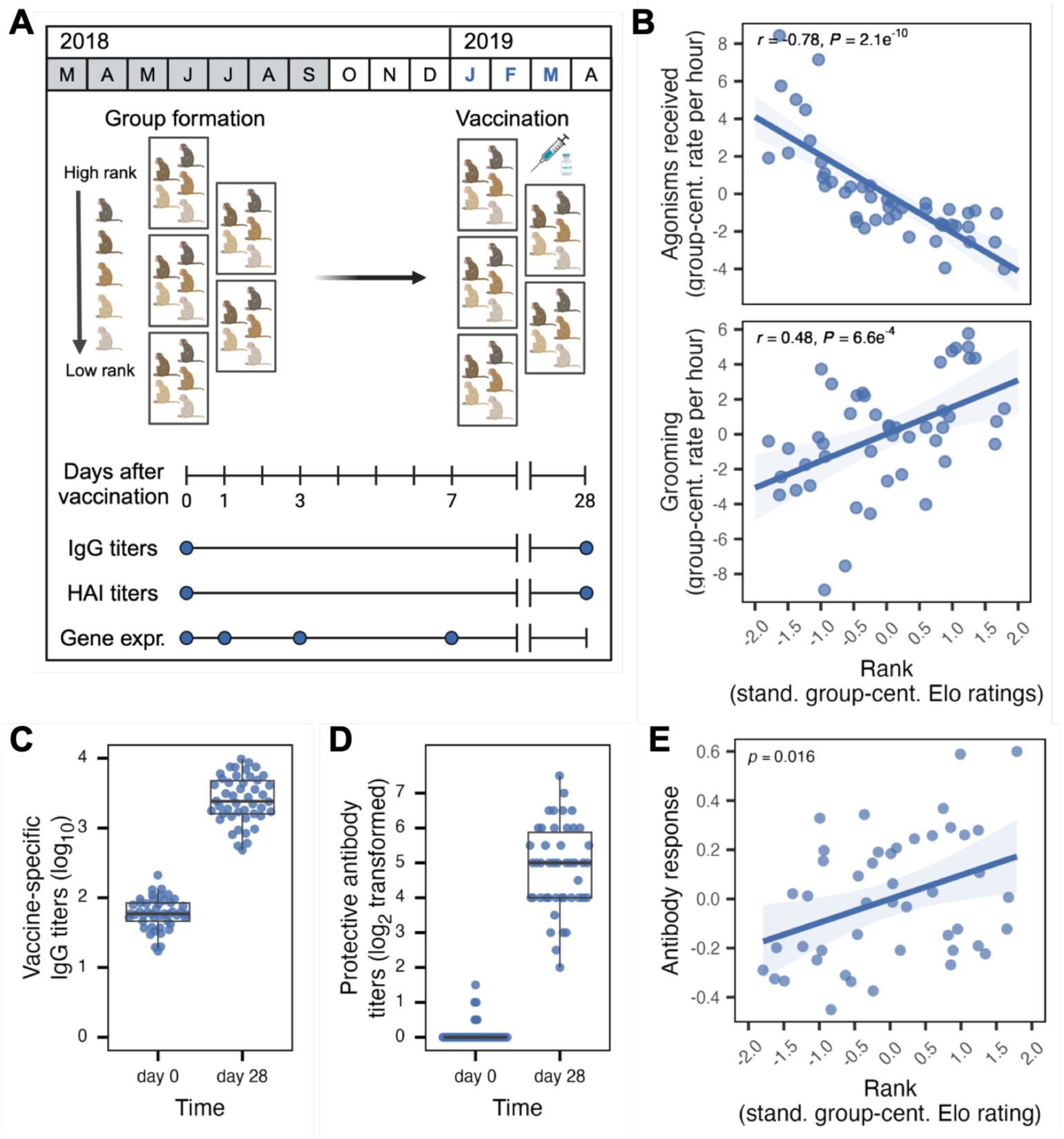
Social status drives social behavior and the antibody response to vaccination. (**A**) Diagram for initial manipulation of social status in rhesus macaques and timeline for group formation, vaccination and sampling, with sampling periods highlighted in blue. (**B**) Relationship between dominance rank (mean-centered Elo rating by group) and group-centered values for agonisms received (top) and overall grooming rates (bottom) based on observations from group formation (March-September 2018) to vaccination date (January-February 2019, see Data S1 for exact dates). High-ranking females received less harassment from their groupmates and engaged in grooming (both given and received) more frequently. (**C**) Titers of vaccine-specific IgG antibodies, measured prior to and 28 days after administration of influenza vaccine to naïve study subjects. Vaccine-specific titers are log_10-_transformed. (**D**) Titers of antibodies protective against influenza-induced hemagglutination, measured prior to and 28 days after administration of influenza vaccine to naïve study subjects. Titers are divided by 5 and log_2_-transformed, following (*128*). (**E**) Effect of dominance rank, measured by Elo rating, on the IgG antibody response (linear mixed model *p* = 0.016). The y-axis shows the partial residuals for antibody titers as a function of dominance rank, adjusted for the contribution of age, body mass, group membership, and kinship.

Next, we administered human pediatric quadrivalent influenza vaccine (Fluarix 2018/2019), containing a mix of two influenza A and two influenza B strains, at 15 µg of influenza virus hemagglutinin/strain. We expected our study subjects to be naïve to influenza exposure prior to inoculation. Consistent with this expectation, we observed a robust antibody response 28 days post-vaccination (paired t-test *p* < 2.2 x 10^-16^; mean fold-increase in vaccine-specific IgG antibody titers relative to the day of vaccination = 43 [95% CI: 34-54]; Fig. 1C and Data S2). Additionally, serum antibodies capable of inhibiting influenza hemagglutination (HAI; a measure of functional humoral immunity) rose from undetectable levels at baseline to a geometric mean HAI titer of 132 by day 28 post-vaccination (Fig. 1D and Data S2). These observations together reflect the universal development of a robust adaptive immune response to the influenza strains contained in the vaccine.

We then investigated our primary question: whether experimentally manipulated social status causally drives variation in the immune response to influenza and, especially, whether high rank confers an advantage. We found that Elo rating at the time of vaccine administration predicts a graded fold-change response in vaccine-specific IgG antibody titers measured 28 days post-vaccination. Higher ranking females (top quartile of Elo ratings) mounted an approximately 1.5-fold (50%) stronger antibody response to vaccination, on average, than lower ranking females (bottom quartile of Elo ratings; linear mixed effects model *p* = 0.016 controlling for age, body mass, group membership, and kinship; Fig. 1E). We observed a qualitatively similar positive correlation between dominance rank and serum antibodies in the HAI assay: females in the top 25% of the Elo hierarchy produced, on average, 1.9-fold higher levels of protective antibodies than those in the bottom 25%. However, when controlling for other covariates, this effect was not statistically significant (*p* = 0.33; Fig. S1).

### Social history influences the secondary “booster” response to influenza vaccination

Unlike the innate immune response, the adaptive immune response is defined by its capacity to influence subsequent exposure to the same pathogen signals. We therefore investigated whether dominance rank at the time of first vaccination continued to predict the antibody response to a secondary booster shot administered approximately one year later. To disentangle the effect of dominance rank history from dominance rank coincident with the booster vaccination, we performed a second social status manipulation by reorganizing similarly ranked members of the initial study groups into new, shared social groups (following (*80*) and (*88*), Fig. 2A).

**Fig. 2.**
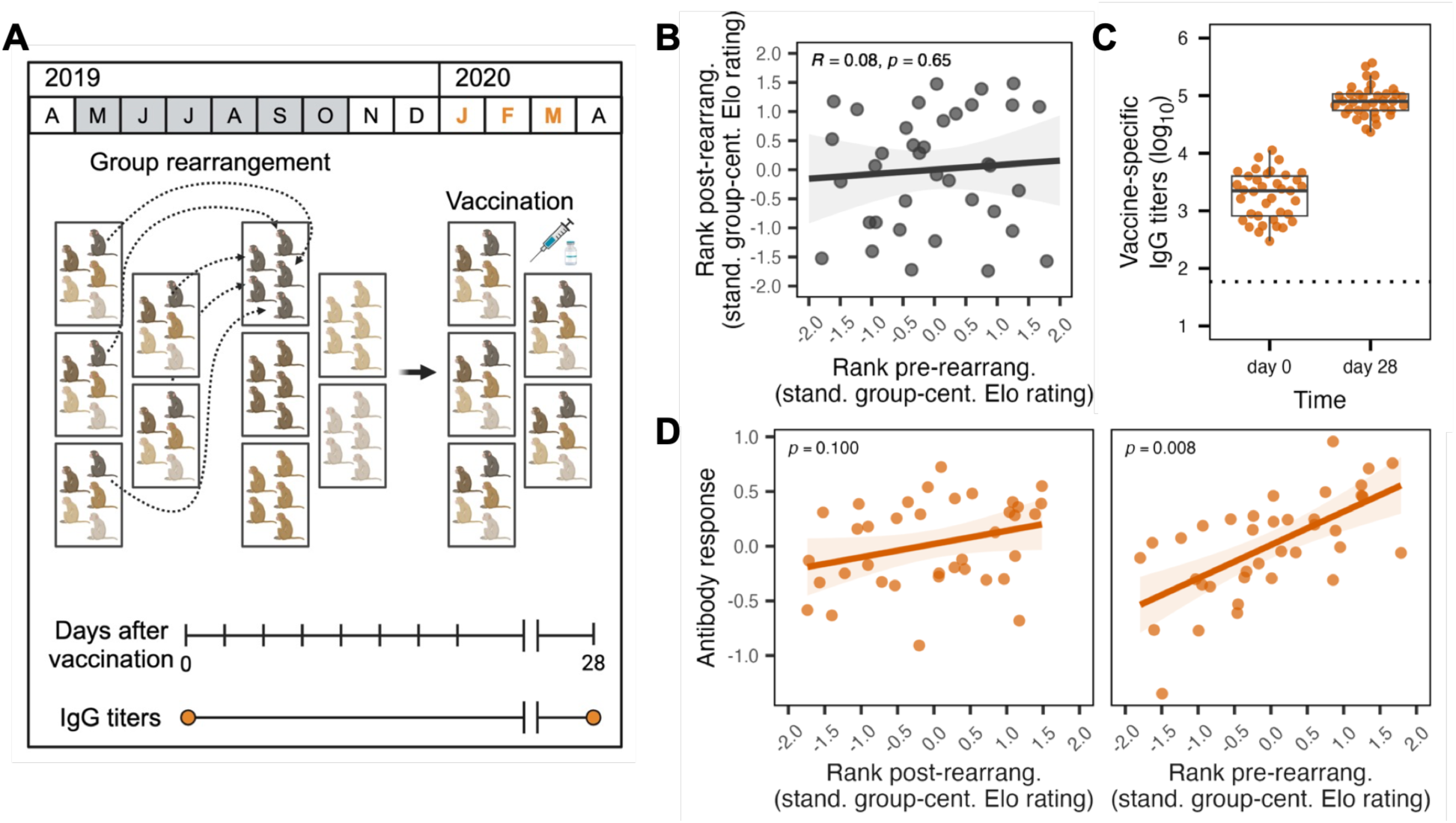
Social history, not current social status, predicts the antibody response to booster vaccination. (**A**) Diagram of the secondary manipulation of social status and timeline for group rearrangement, booster vaccination, and sampling, with sampling periods highlighted in orange. (**B**) Dominance rank (Elo rating) is uncorrelated between the first and second rounds of vaccine administration, as expected based on the study design. (**C**) Titers of vaccine-specific IgG antibodies measured prior to and 28 days after administration of the second influenza vaccine. For reference, the dotted line denotes baseline levels of IgG antibodies prior to the first immunization. (**D**) Effect of dominance rank (Elo rating) on the IgG antibody response to the second (booster) vaccination. Both plots show partial residuals for antibody titers as a function of dominance rank, adjusted for the contribution of age, body mass, group membership, and kinship. There is no effect of Elo rating measured at the time of booster vaccination on the secondary response to vaccination (left; linear mixed model *p* = 0.100). However, Elo rating at the time of first vaccination significantly predicts the IgG antibody response a year later (*p* = 0.008).

This approach generated Elo ratings that were completely uncorrelated within-subject before and after the second status manipulation (Pearson’s *r* = 0.08, *p* = 0.65, Fig. 2B). Elo rating post-manipulation was also predicted by order of introduction (Pearson’s *r* = −0.42, *p* = 3.6 x 10^-3^). After groups stabilized, we administered a second quadrivalent influenza vaccination, which was identical to the 2018/2019 vaccine except for a single influenza A strain (see Materials and Methods). 28 days after administration, the booster shot produced an average 44-fold increase (95% CI: 34-57) in vaccine-specific IgG titers relative to the day the animals received the shot (paired t-test *p* < 2.2 x 10^-16^ for day 0 to day 28 titers). This increase was similar in magnitude to the original response, but absolute antibody titers were substantially higher than after the first vaccine because all animals retained vaccine-specific antibodies from the initial vaccine administration (Fig. 2C).

Unlike our findings in the first phase of the study, dominance rank at the time of the booster administration did not significantly predict the magnitude of the antibody response (linear mixed effects model *p* = 0.1, controlling for age, body mass, group, and kinship; Fig. 2D, left panel). In contrast, Elo rating at the time of the initial vaccination – when animals were immunologically naïve – remained a significant predictor of the secondary response (*p* = 0.008; Fig. 2D, right panel). These results are consistent with the idea that individuals with higher social status at the time of first antigen exposure develop a memory B cell pool that is more effective at mounting a recall response upon re-exposure.

### Social status drives differential expression of adaptive immune genes that are specifically active late in the response to vaccination

We next asked whether social status effects on the response to vaccination are connected to the frequently reported link between social status and gene expression levels in peripheral immune cells (*79*, *80*, *88*, *95*, *98*). To investigate this possibility, we characterized both the gene expression signature of social status and the gene expression response to the influenza vaccine in the week after the first vaccine administration, in peripheral blood mononuclear cells (PBMC) (n = 43 individuals, see Fig. S2 and Materials and Methods, sampled pre-vaccination [day 0] and days 1, 3, and 7 post-vaccination).

We identified 4,782 unique genes that were differentially expressed, relative to the day 0 (pre-vaccination) sample, at one or more time points post-vaccination (false discovery rate [FDR] = 1%, controlling for age and body mass; Data S3). Although more genes were shared across time points than expected by chance, many cases were specific to the early versus late immune response to influenza antigens. Unsupervised clustering into six possible trajectories of response (Fig. S3 and Data S4) highlighted the primary role of innate immunity on day 1: genes that were transiently up-regulated at day 1, but that returned to baseline levels by day 3 (Cluster 1), were enriched for interferon alpha and gamma signaling, neutrophil degranulation, leukocyte migration and the regulation of viral genome regulation (all *p_adj_* < 1.0 x 10^-3^; Fig. 3A left panel, Data S5). In contrast, day 7 reflects the activation of adaptive immune pathways and the downregulation of early-response inflammatory pathways. Genes specifically upregulated by day 7, but not at earlier time points (Cluster 2), are enriched for antigen cross-presentation and B and T cell receptor signaling (Fig. 3A middle panel, Data S5). Genes that were down-regulated from day 3 to day 7 (Cluster 3) are enriched for the inflammatory response and interleukin signaling (all *p_adj_* < 2.4 x 10^-3^; Fig. 3A right panel, Data S5). The other three clusters exhibited no discernable relationship with annotated immune pathways or processes.

**Fig. 3.**
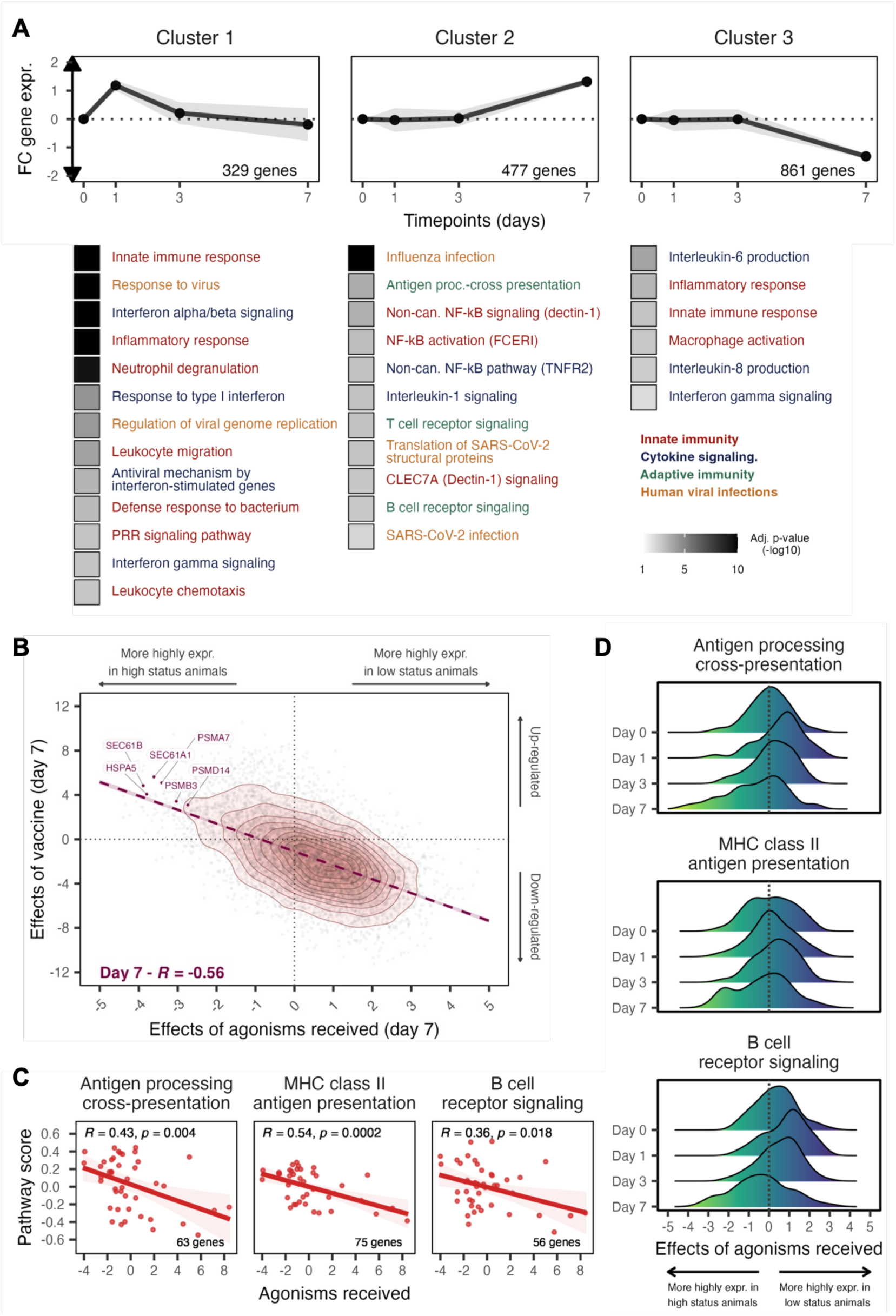
Concordance between gene expression responses to influenza vaccination and gene expression responses to status-related agonistic behavior. (**A**) Temporal dynamics of gene expression among gene clusters (top) and over-representation enrichment analysis for Reactome pathways and Gene Ontology terms among those clusters (bottom). Immune terms are colored according to broad immune categories. (**B**) Correlation of gene-level effect sizes (standardized betas) between the vaccine response and agonisms received at day 7 following vaccination. Examples of genes that are both significantly affected by influenza antigen exposure (genes upregulated following vaccination, FDR 1%) and social behavior (genes more highly expressed in females rarely targeted by their groupmates, FDR 20%) are labeled. (**C**) Pathway scores for three gene sets related to adaptive immunity, based on mean day 7 gene expression across genes in each pathway, are correlated with the rate of agonisms received. Scores reflect the relative enrichment in genes from a pathway in a given sample. (**D**) Effects of agonisms received on the expression of genes annotated in adaptive immunity pathways are not consistent over time, but include the clearest outliers on Day 7.

To identify genes affected by dominance rank, we focused on modeling the rate of agonisms received (e.g., threats, spatial displacements) by each individual. More agonistic behavior is directed towards low-ranking female macaques than high-ranking macaques (Fig. 1B), and previous work suggests that this experience mediates social status-related gene expression levels (*85*). While the signature of dominance rank is weaker in this data set than in previous studies (*79*, *80*, *88*), the day 7 signature (n = 733 agonism-associated genes, FDR = 20%, Data S6) is strikingly concordant with the gene expression response to vaccination at the same time point. Gene-level effect sizes for these two predictor variables were more highly correlated on day 7 (Pearson’s *r* = −0.56, *p* < 1.0 x 10^-10^, Fig. 3B) than on days 1 or 3 (day 1: *r* = - 0.27, *p* < 1.0 x 10^-10^; day 3: *r* = −0.27, *p* < 1.0 x 10^-10^; Fig. S4). Consequently, on day 7, genes that were upregulated in response to vaccination overlapped with genes that were more highly expressed in females who were rarely targeted by agonistic behavior (log_2_(OR) = 3.7, *p* < 1.0 x 10^-10^). These females, who all tended to be high ranking, exhibited mean increased expression of genes involved in MHC class I antigen cross-presentation (Pearson’s *r* = 0.43, *p* = 0.004, Fig. 3C, left panel, Data S7), MHC class II antigen presentation (*r* = 0.54, *p* = 0.0002, Fig. 3C, middle panel, Data S7), and B cell receptor signaling (*r* = 0.36, *p* = 0.018, Fig. 3C, right panel, Data S7), relative to low ranking females. Notably, many of these genes are either uncorrelated with agonism rates or show the opposite effect earlier in the time course (Fig. 3D).

### High status females upregulate genes that predict stronger vaccine-specific IgG antibody responses

Our findings above indicate that experimentally manipulated social status is sufficient to drive both differences in the antibody response to vaccination and differences in the regulation of adaptive immune pathways that are expected to partially undergird this response. To connect these two observations, we therefore asked whether cases in which gene expression levels strongly predict IgG antibody titers 28 days after vaccination – the last time point we collected data post-vaccination and a major predictor of lasting immunity (*99*, *100*) – are also polarized by social status.

The fold-change increase in vaccine-specific IgG antibodies from day 0 to day 28 was most strongly predicted by the changes in gene expression observed at day 1 after immunization, when the innate immune response to vaccination is most pronounced. We identified 1,882 Ab-associated genes (FDR < 10%), 1,438 (75%) of which were negatively correlated with the antibody response (hereafter referred to as “negative predictors”) and 444 (25%) of which were positively correlated (hereafter referred to as “positive predictors”). Positive predictors tended to fall in innate pro-inflammatory immune categories such as regulation of granulocyte activation, Toll-like receptor cascades, and interferon gamma signaling (all *p_adj_* < 4.0 x 10^-3^; Data S8). Gene Set Variation Analysis (GSVA) pathway enrichment scores (*101*), which capture a sample-specific estimate of whether genes from a given pathway are expressed higher or lower as a group relative to the rest of that sample’s transcriptome, strongly predicted fold-change differences in antibody titers at day 28 (Fig. 4A). Notably, nearly all of the strongly upregulated, positively predictive genes were interferon-stimulated genes (ISGs) with known antiviral functions (e.g., *OASL*, *APOBEC3A*, *IFIT1*, and *MX1*; Fig. S5) that also appear in gene expression signatures of the antibody response to influenza vaccination in humans (24–100% overlap; log₂(OR) = 1.47 to 4.67, *p_adj_* = 0.002 to 6.0 × 10⁻²⁹; Fig. 4B) (*102–106*).

**Fig. 4.**
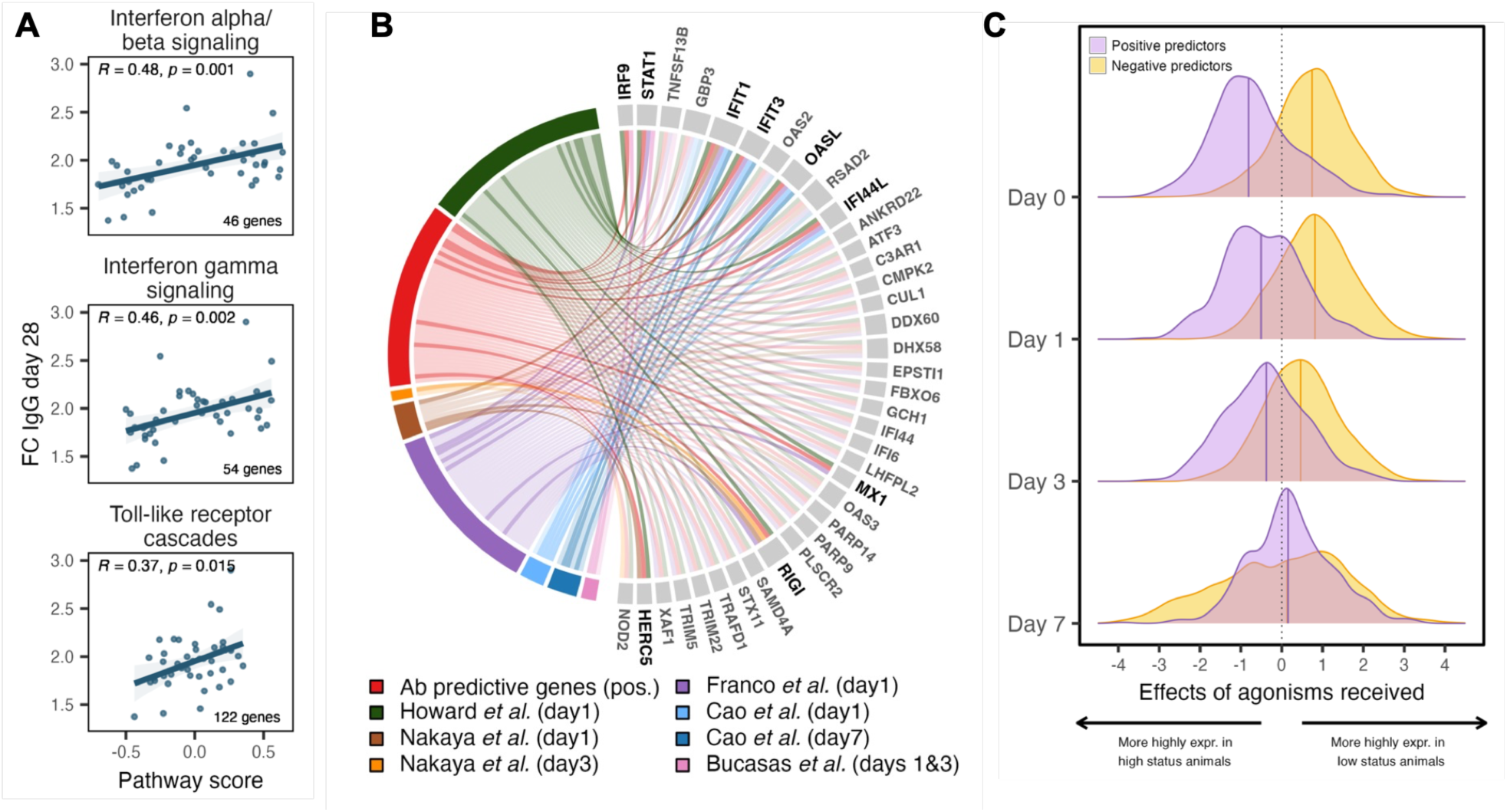
Gene expression predictors of the antibody response to influenza vaccination are also polarized by social status-related social behavior. (**A**) Pathway scores for the interferon response and TLR signaling pathways, based on day 1 differential gene expression, predict the fold-change IgG antibody response at day 28. (**B**) Positive antibody predictive genes identified here overlap with those reported in human influenza vaccination studies (*102–106*). The circos plot shows Ab-predictive genes that are also found in at least 2 other data sets in humans, with immune-related genes highlighted in bold. (**C**) Distribution of effect sizes for the relationship between agonisms received and gene expression for positive predictors (purple) and negative predictors (yellow) of the antibody response. Differences between the two distributions were quantified using the Wilcoxon rank-sum test (BH-adjusted). Early timepoints show strong distributional separation that diminishes by day 7.

Finally, we asked whether high status females showed elevated expression of positive predictors and reduced expression of negative predictors, consistent with their stronger antibody responses. We found that the effects of agonisms received on gene expression differed in sign between the two gene sets. Among positive predictors of antibody responses, effect sizes were predominantly negative, indicating higher expression in high-ranking females. In contrast, among negative predictors, effect sizes in the agonism data were predominantly positive, indicating higher expression in low-ranking females (Fig. 4C, Wilcoxon rank-sum test comparing the distribution of effect sizes between positive and negative predictors, *p_adj_* < 2.2 × 10⁻¹⁶ for days 0, 1, and 3; *p_adj_* = 0.07 for day 7). This status-linked polarization was strongest at baseline and day 1, when gene expression is most informative of later antibody levels, and diminished by day 3 and day 7 (Fig. 4C). Accordingly, the average expression of positive predictors was strongly correlated with social status, reinforcing the idea that immune transcriptional responses that support lasting antibody production are shaped by social experience.

## Discussion

In the current study, we demonstrate that the immune response to influenza vaccination in female rhesus macaques is influenced by their social status at the time of first vaccination. Given that low status rhesus macaques often exhibit markers of chronic stress, including high glucocorticoid levels and glucocorticoid resistance (*78*, *83*, *84*), our findings support the idea that social gradients in infectious disease phenotypes arise in part through the immune dysregulation caused by chronic social stress. Our results therefore provide experimental evidence for a causal pathway that may also be at play for associations between chronic psychosocial stress, viral infection, and vaccine efficacy in humans (*56*, *107*), including in response to the influenza vaccine (*50*, *53*).

However, our findings indicate that this relationship has an important experiential component: social status upon initial exposure to influenza antigens affects both the initial antibody response and the efficacy of a subsequent booster, but we detected no effect of social status at the time of the booster. Our results are consistent with growing evidence that initial immune responses in antigen-naïve individuals can predict the magnitude of later booster responses. For example, antibody levels after the first dose of a COVID-19 vaccine have been shown to correlate with post-boost titers in both elderly populations (*108*) and adult healthcare workers (*109*), particularly for mRNA-based vaccines (*110*). For both COVID-19 and influenza, sustained immunity depends on the maintenance of specialized memory cells generated during the primary response, which increase the speed and strength of secondary immune responses (*111*). Our data therefore provide interpretative clarity to observations in humans, which show that impairments in antibody responses in stressed caregivers persist across flu seasons, even when the source of chronic stress abates (*50*). Together, these results suggest that studies of the relationship between social disadvantage and infectious disease can fail to identify a true relationship, if the trait of interest depends on adaptive immunity and if data about social conditions at earlier exposure times are unavailable.

Our result contrasts with studies focused on the innate immune response, which appears to be much more plastic in response to changes in prevailing social conditions (*112–116*), including at the level of gene expression (*80*, *88*). However, our analysis also shows how gene expression signatures of the social environment connect to slower-developing adaptive immune responses to pathogen stimuli. We find that the signature of social status is most pronounced a week after vaccine administration, the period post-exposure when the immune system transitions from a primarily innate to a primarily adaptive response. As in humans (*103*, *117*, *118*), this transition is marked by a shift from an early, transient activation of interferon signaling and pro-inflammatory pathways to the later induction of lymphocyte receptor signaling and antigen processing and presentation pathways. At that time, genes in both of these adaptive immune pathways are differentially regulated as a function of social status. For example, genes encoding subunits of the MHC I peptide-loading complex and the channel-forming translocon complex are more highly expressed in high status females. Both complexes are critical for antigen loading and export from the endoplasmic reticulum to the cell surface (*119*, *120*) and for the transport of antigen peptides and MHC molecules between intracellular compartments (*121*). Improved activation of T cells in high status females could, in turn, lead to more B cell proliferation and immunoglobulin production (*122*), which would explain the stronger antibody responses observed at day 28.

Our findings may also help clarify why associations between social conditions and antibody levels in humans differ across pathogens. Our results are consistent with pathogen-dependent associations between SES and antibody levels following vaccination in humans (*123*). Higher SES was associated with increased IgG titers against measles and rubella, presumably reflecting the primary response mounted during childhood exposure in naïve individuals, as these viruses typically cause disease in childhood and are cleared. In contrast, IgG titers against pneumococcus and CMV, which usually cause recurrent or persistent infections, as well as meningitis C, were higher among low-SES subjects, presumably due to higher exposure and/or pathogen burden in disadvantaged individuals. Our experimental design isolates immune programming from variation in exposure and demonstrates the importance of the social environment specifically at first antigen encounter. We therefore speculate that social gradients in antibody levels will depend on whether a given pathogen primarily reflects primary immune imprinting or ongoing antigenic stimulation.

Together, our findings highlight the role of the social environment as a major driver of antiviral immune responses, particularly during initial pathogen exposure in naïve individuals. They suggest that social gradients in infectious disease therefore not only reflect important differences in preventative care, exposure, and treatment (*1*, *44*, *46*, *48*), but also physiological differences at the cellular and sub-cellular levels that influence how individuals respond to infectious agents. Notably, widespread evidence indicates that social disadvantage and chronic social stress are associated with increased inflammation—a process that can be tissue-damaging, but also protective against pathogens when deployed in the right context (*124*). Our results support this idea in part: early activation of inflammatory genes predicted a better vaccine response. In addition, they point to temporally specific activation of these pathways, in a pattern linked to relative social advantage, as a key driver of effective immunity against influenza.

## Materials and Methods

### Study Animals and Experimental Design

Study subjects in phase 1 were 47 adult female rhesus macaques (ages 5.6 – 25.8 years; mean age at vaccination = 14.1 years ± 4.81 s.d.) housed in groups of five females each at the Emory National Primate Research Center (ENPRC) Field Station (Data S1). The ENPRC is fully accredited by AAALAC International and all data were obtained in accordance with Institutional Animal Care and Use Committee (IACUC) protocols approved by Emory University (PROTO201700750). All study subjects, except for three individuals who originated from external animal colonies, were born in the ENPRC breeding colony. Study subjects were housed under two conditions: 24 females were kept in specific-pathogen-free (SPF) conditions, where animals are confirmed seronegative for Simian Immunodeficiency Virus (SIV), Simian T-Lymphotropic Virus (STLV), Simian Type D Retrovirus (SRV), and Cercopithecine Herpes B (B-virus) in annual testing. 23 females were housed in non-SPF conditions. We therefore performed additional tests for the presence of SIV, STLV, SRV and B virus in the non-SPF animals, resulting in three positive STLV cases and 11 positive B-virus cases (none for the other two pathogens). However, these positive cases were random with respect to social status at the time of sampling (Wilcoxon rank-sum test, *p* = 0.53 for STLV and *p* = 0.53 for BV).

All ten social groups in this analysis were formed in March – October 2018 and rapidly formed the stable, linear, and regularly enforced dominance hierarchies typical of female rhesus macaques (*125*). Group construction followed a well-established protocol (*78*) in which 4 – 5 females were sequentially introduced into indoor-outdoor run housing (144 square feet per run) over the course of several weeks and kept on unrestricted access to a normal low-fat, high-fiber nonhuman primate diet. Observational behavioral data collection began on each social group once it was complete (i.e., the final female macaque was introduced into that group). Vaccination occurred 19-46 weeks after group completion and the start of behavioral observations (median = 33 weeks). Samples collected for antibody, HAI assay, and gene expression analysis were collected relative to each individual’s vaccination date, as shown in Fig. 1A.

Nine of the individuals in phase 1 were no longer in the study cohort by the time we performed “booster” vaccinations in phase 2. Five of these animals were replaced with new individuals, to retain similar social group sizes in phase 2 compared to phase 1. Replacement animals were included in behavioral analysis (e.g., to define rank and to measure agonism/grooming rates) but excluded from gene expression (conducted in Phase 1 only) and antibody response analyses because their history of antigen exposure differed from those of our primary study subjects (see sections below). Consequently, we retained 38 individuals for antibody analysis in phase 2, all of whom had matched data from phase 1. Social groups in phase 2 were formed following the same serial introduction approach as in phase 1: in this case, to shuffle ranks among the animals in the data set, we grouped females of the same or adjacent ordinal ranks in phase 1 into the same group in phase 2. Consequently, within-individual rank values across phases are uncorrelated (Pearson’s r = 0.08, *p* = 0.65, Fig. 2B), allowing us to disentangle causal effects of social status at the time of sampling from historical social experience. We excluded one female from the analysis in phase 1 because she was a major outlier in the behavioral data (see *Behavioral Measures* section). Our final sample sizes for antibody analyses were therefore 46 individuals in phase 1 and 37 individuals in phase 2 (Data S1).

### Behavioral Measures

To monitor dominance rank, we collected behavioral data once per week per group using group-level focal observations (278 total hours of observation, 180.5 hours in phase 1 and 97.5 hours in phase 2). We assigned dominance rank by calculating Elo ratings (*97*, *126*), a continuous measure of rank in which animals with higher Elo ratings have higher social status, and ratings are adjusted following any dyadic interaction in which a winner or loser can be determined. Wins increase the winner’s Elo rating and losses decrease the loser’s Elo rating by an amount that is proportional to the expected outcome of the interaction. For instance, if an individual with a high Elo rating wins an interaction with an individual with a low Elo rating, an expected outcome based on their scores, their Elo ratings change little. If the outcome is flipped, however, the scores change more to more accurately capture the social hierarchy.

We calculated Elo ratings for each social group using all agonistic interactions that occurred after the final female was introduced into a social group. The initial Elo rating for each female was set to a value of 1,000, and the baseline number of points a female could potentially gain or lose during a dominance interaction (k) was set to 100. For each interaction, k was weighted by the expected probability of an individual winning or losing, based on a logistic function that was updated after each dominance interaction. The animals’ order of introduction to their group enclosure predicted subsequent dominance rank at the time of vaccination (Pearson’s *r* = −0.51, *p* = 1.9 x 10^-4^, n = 49 females for phase 1 and *r* = −0.42, *p* = 3.6 x 10^-3^, n = 46 for phase 2). Dominance structure was consistent with moderate-to-high linearity (mean h′ = 0.99 for first phase and mean h′ = 0.975 for second phase), as expected for this sex-species combination.

To test whether Elo ratings predicted social interactions, as expected, we assessed rates of received harassment and grooming. Received harassment was defined as the per-hour rate of received agonisms directed from a focal female’s groupmates towards her, and grooming rates were defined as the amount of time, in minutes per hour, the focal female was engaged in either giving or receiving grooming from her groupmates. We mean-centered these values across groups to 0, such that our measures of received harassment and grooming are relative to the group mean (i.e., reflect whether a female was targeted more often or less often than the typical value for her group). This decision removes potential technical differences between groups that could emerge, for example, if grooming or agonisms were measured more frequently in some groups than others because of observer effects. However, it makes the implicit assumption that the *relative* rate of received harassment or grooming is more salient to our outcome variables than the *absolute* rate in which animals experience these types of interaction.

Overall, the behavioral data for the study sample followed expected patterns for female rhesus macaque social hierarchies. Grooming was more frequent for closely ranked individuals, based on the absolute difference between Elo ratings (generalized linear mixed model with log link for total grooming duration (in minutes): β = −0.712, *p* = 9.94 x 10^-6^ in phase1; β = −0.665, *p* = 1.17 x 10^-5^ for phase 2, controlling for actor and recipient identity and dyad identity) and biased up the hierarchy (Fig. 1B bottom panel). Elo ratings remained highly stable across each phase (S = 0.997 for first phase and S = 0.997 for second phase). However, a single individual, “DV2J”, had behavioral values that were conspicuous outliers in both the grooming and agonism data, leading to an unusually low Elo score. As this individual’s social experience fell well beyond the range of typical experiences for female macaques in this study paradigm, we removed her from the data set before further analysis, leaving the total number of analyzed individuals at 46. Any behavioral metrics calculated for her groupmates (Elo interactions with DV2J, grooming or agonisms involving DV2J) were left in the dataset, however the values for DV2J were dropped and behavioral values for members of her group were mean-centered without considering values for DV2J.

### Vaccine Administration

We immunized a cohort of rhesus macaques with human quadrivalent influenza vaccine (Fluarix, GlaxoSmithKline Biologicals). Vaccine doses were administered twice, with an approximately one-year interval between doses: the first dose after initial group formation (phase 1, January 2019) and the second dose following group rearrangement (phase 2, January 2020).

In phase 1, 47 female rhesus macaques were injected intramuscularly with 500 µl of reconstituted influenza vaccine (2018/2019). This vaccine was composed of four influenza strains: two A (A/Michigan/45/2015 and A/Singapore/INFIMH-16-0019/2016) and two B (B/Colorado/06/2017-like virus and B/Phuket/3073/2013-like virus) strains and contained 15 µg of hemagglutinin/strain. The vaccine was administered together with a commonly used adjuvant (Addavax, Addgene) to ensure proper immune response in naïve animals. All vaccinations occurred within the span of one month (January to February 2019; see Data S1).

After group rearrangement (phase 2), the 37 animals who continued in the study from Phase 1 were administered a second dose of human quadrivalent influenza vaccine (2019/2020), following the same procedures performed in Phase 1. This vaccine differed from that administered in phase 1 by a single influenza A strain (A/Kansas/14/2017 instead of A/Singapore/INFIMH-16-0019/2016).

In Phase 2, we also included five new individuals to replace those removed from the study prior to the start of phase 2. However, because these animals only received one dose of the vaccine, they were included to calculate Elo ratings and behavioral metrics but not included in the antibody response analysis (see Study Animals and Experimental Design section above and Data S1).

### Blood Sample Collection

In phase 1, we collected peripheral blood samples from all animals (n=46) before immunization (“day 0” phase 1) and at four additional timepoints following vaccination (days 1, 3, 7 and 28 in phase 1). From blood drawn on days 0, 1, 3 and 7, we isolated PBMCs for RNA extraction and sequencing. Briefly, 3 mL of blood collected in Vacutainer Mononuclear Cell Preparation Tubes (CPT) with sodium citrate (BD) were centrifuged at 1800 g for 30 minutes to promote separation of cellular and plasma fractions. 500 µl of plasma were collected from the upper fraction, transferred to 1.5 mL cryovials, and stored at −80 °C. To isolate PBMCs, we transferred the remaining fraction to a new tube, washed it with PBS 1X, and lysed the cells in 350 µL of RLT buffer (Qiagen) containing 2-mercaptoethanol. Cell lysates were stored at −80 °C until RNA extraction. In addition to PBMCs, we collected plasma on day 0 for antibody assays.

To measure the 28 day antibody response, we also isolated plasma from blood samples collected 28 days after vaccination, following a similar protocol. Blood samples (2 mL) collected in Vacutainer Acid Citrate Dextrose (ACD) tubes (BD) were centrifuged at 1500 for 10 minutes. After separation of blood fractions, 500 µL of plasma were transferred to 1.5 mL cryovials and stored at −80 °C.

In phase 2, we collected plasma samples from individuals prior to immunization (day 0, phase 2) and 28 days after the second immunization (day 28, phase 2) following the protocol described above for phase 1.

### ELISA assays for vaccine-specific IgG quantification

We performed enzyme-linked immunosorbent assays (ELISAs) to quantify the levels of vaccine-specific IgG antibodies in the plasma of the rhesus macaques in our study. Prior to their use in immunological assays, plasma samples were subjected to a heat inactivation step by heating them to 56 °C for 45 minutes to neutralize any viral contaminants that could have been present.

Standard 96-well microplates (Costar 3369) were coated with 50 µL of a 20-fold dilution in PBS of the same quadrivalent influenza vaccine administered to the animals (Fluarix), followed by overnight incubation at 4 °C. Coated plates were then washed three times with wash buffer (PBS 0.05% Tween) and blocked with blocking solution (PBS 20% FBS, 150 µL/well). After 1 hour incubation at 37 °C, plates were washed again three times with wash buffer. We prepared dilutions of inactivated plasma in PBS, including a 10-fold initial dilution and then seven serial 4-fold dilutions. 50 µL of plasma dilutions or PBS control were added to the coated ELISA plates (two replicates per sample) and incubated for 1 hour at 37 °C. After three washes with wash buffer, 75 µL of HRP-conjugated anti-rhesus macaque IgG antibody (Sigma, 1000-fold dilution in PBS) were added to the ELISA plates. After a one hour incubation at 37 °C and three washes, developing reagent (Super AquaBlue ELISA substrate, EBioscience) was added to the plates (100 µL/well), followed by a final room temperature incubation in the dark for exactly 20 minutes. Color produced by the enzymatic reaction was evaluated by reading the absorbance at 405 nm in a microplate reader (Multiskan GO, Thermo Scientific). All plasma samples were measured twice in independent ELISA assays.

IgG titers were measured from plasma samples collected at two timepoints: prior to (day 0) and 28 days after vaccine administration, for both phases 1 and 2. In phase 1, samples were tested in batches, where each batch corresponded to a single social group (up to five individuals per ELISA plate), with all tests conducted on the same day. To ensure direct comparability between titers measured in phase 2 and those from phase 1, we also repeated measurements using plasma samples collected from naïve animals (day 0 phase 1) at the same time we tested samples from phase 2. Thus, samples from these three timepoints (day 0 phase 1, day 0 phase 2 and day 28 phase 2) were randomized, with groups of five samples tested per ELISA plate.

We determined endpoint titers for plasma vaccine-specific IgG using a sigmoid dose response curve (GraphPad Prism, version 9). Titers were defined as the reciprocal of the highest dilution of test plasma that gave an absorbance above the cutoff of the average, plus eight standard deviations of the values obtained from negative (PBS) control wells, see Data S2.

### Hemagglutination inhibition assays

To assess the functional antibody response to vaccination we performed hemagglutination inhibition assays (HAI) with A/Michigan/45/2015 (H1N1)pdm09-like influenza virus, a viral strain present in both vaccines administered during phase 1 and 2. The HAI quantifies serum antibodies that bind influenza virus hemagglutinin, preventing agglutination of red blood cells (RBC) (*127*).

Serum samples were treated with receptor destroying enzyme (RDEII) for 18 h at 37 °C, inactivated at 56 °C for 30-40 min, and diluted 10-fold in PBS. Sera were then transferred in duplicate to V-shaped 96 well plates and subjected to further two-fold serial dilutions in PBS. 4 hemagglutination units (HAU) (25 µL) of pre-titrated influenza virus were added to the serum dilutions and the plates were then incubated for 1 h at room temperature to allow antibody-virus neutralization. 50 µL of turkey RBC (0.5% concentration) were added to each well, followed by incubation at room temperature for 45 min. Positive and negative controls were also included in each plate, i.e., wells containing both influenza virus and turkey RBC but no serum or only RBC. We quantified the titers of protective antibodies against influenza virus for phase 1, at baseline (day 0) and 28 days after immunization with flu vaccine. We tested one social group per plate, with all individuals tested twice in independent assays. HAI titers were determined as the reciprocal of the highest dilution of serum that inhibited hemagglutination of RBC, see Data S2.

### Statistical analyses of antibody response

Vaccine-specific IgG levels (quantified by ELISA) and protective antibody titers (measured by hemagglutination inhibition (HAI) assays) are presented as geometric mean titers from two independent assays. HAI titers from phase 1 were divided by 5 and transformed using a binary logarithmic scale (*128*). On this scale, because the serum was initially diluted 10-fold, undetectable antibody titers (less than 10) are represented as 0, a titer of 10 as 1, a titer of 20 as 2, and so forth. Vaccine-specific IgG titers for phase 1 and 2, determined by ELISA, were log_10_-transformed.

To test whether social rank affected antibody responses to flu vaccination, we fitted a mixed effects linear regression model (R function *lmekin*, package coxme (*129*)), with fold change of antibody titers from day 28 (endpoint) to day 0 (pre-vaccination) as the response variable and dominance rank, represented by group-centered Elo rating, as the predictor variable of interest. We also included social group, mean-centered body mass, and mean-centered age of the animals as fixed effect covariates and accounted for relatedness between individuals by including a kinship coefficient matrix in the following model:

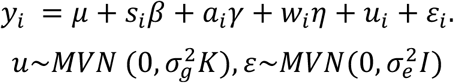

where *i* indexes the study subject and *y* is the *n* x 1 vector of log-transformed fold changes in vaccine-specific IgG titers from day 28 to day 0; *s* is an *n* x 1 vector of Elo ratings and β the effect size of Elo scores on antibody titers. The vectors *a* and *w* are covariates with associated fixed effects γ and η for age and body mass at Day 0, respectively. The random effects term 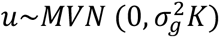 controls for kinship and other sources of genetic structure, where K is an *n × n* matrix of pairwise relatedness estimates derived from genotypes at 62,648 single nucleotide variants (SNVs) (see *Genotyping* section below). Residual errors are modeled as 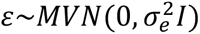, where *I* is the identity matrix and 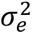 captures the environmental variance component.

We fit a separate model, with identical structure, for each of the two phases. Fold change for phase 2 was adjusted for baseline titers using Beyer’s correction (*128*), as the animals were no longer naïve for influenza. We also calculated fold change of HAI titers for phase 1 as the difference between log-transformed titers before and after vaccination.

### RNA-Seq library preparation, read mapping, and batch correction

We isolated total RNA from cell lysates using the miRNAeasy kit (Qiagen), following the manufacturer’s instructions. We quantified the concentration of the extracted RNA using a Qubit instrument (Thermo Fisher Scientific), assessed RNA integrity on an Agilent Bioanalyzer (Agilent RNA Nano chip), and confirmed the purity of the RNA extracts by measuring 260/280 and 260/230 ratios on a Nanodrop instrument (Thermo Fisher Scientific). For one individual (Iv8), the RNA yield was consistently low and insufficient for sequencing. We therefore excluded Iv8 from the analysis, resulting in a total of 172 samples across the four vaccine timepoints and from 45 unique individuals.

RNA libraries were prepared using 300 ng of total RNA using the Illumina Stranded mRNA Prep kit (Illumina) according to the manufacturer’s protocol. Libraries were sequenced with single-end 100 bp reads on an NovaSeq X (Illumina) to an average depth of 15 million reads per sample. The resulting reads were trimmed with trimmomatic (*130*) to remove adapter sequence and trailing bases with a quality score less than 20. The trimmed sequence data were then mapped to the rhesus macaque genome (RheMac8) using the STAR v2.7.10ac (*131*) two-pass method. Read counts were generated for each gene using HTCseq v2.0 (*132*) based on overlap of any part of a read with an annotated gene exon. The final set of read counts were then analyzed in R v4.1.2 (*133*). We removed samples from two individuals, RIe6 and RAi10, because for all time points, samples from these individuals were clear outliers in a principal components analysis of the gene expression data (Fig. S2), even after controlling for group, batch, and other technical effects (see below). We therefore retained a total of 43 unique individuals in the data set.

Before data analysis and modeling, we filtered out any genes that were either not detectable or expressed at a low level in our samples. We set a minimum threshold of mean log(CPM) > 3 at any of the three timepoints, resulting in a set of 8,254 analyzed genes. We normalized the gene count matrix using the function *voom* (R package limma, v3.60.4 (*134*)). To account for potential batch effects and technical artifacts, we regressed out the social group of the animal, sequencing run lane, total gene counts, and unique genes identified in each sample using a linear model in R. Since vaccinations and samplings were conducted by social group, controlling for social group identity effectively mitigates many technical batch effects associated with the sample collection process. Additionally, adjusting for sequencing lane, total gene counts, and unique genes addresses artifacts related to individual library quality. All subsequent analyses were based on the residuals from this model.

### Genotyping

We used genotype data to confirm sample identity and control for genetic relatedness between individuals. Briefly, RNA-seq data were generated from cryopreserved PBMCs collected from all individuals prior to the start of the vaccination experiments. We then joint genotyped the samples using the GATK Best Practices for RNA-seq data and the following hard filters (following (*80*): Quality Score > 100.0; QD < 2.0; MQ < 35.0; FS > 60.0; HaplotypeScore > 13.0; MQRankSum < −12.5; ReadPosRankSum < −8.0). The resulting SNPs were thinned to be >10 kb apart (n = 62,648 SNVs) and kinship coefficients were estimated with lcMLkin (*135*). While average pairwise genetic relatedness among individuals was low (0.014), several closely related individuals (r>0.125) were included in the data set. These dyads were never included in the same experimental group in either of the two phases.

### Modeling Effects on Gene Expression

To identify genes that were significantly associated with agonisms received, we used a linear mixed effects model that controls for relatedness within the sample, using the R package EMMREML (*136*). The response variable in this model was batch-corrected gene expression values after controlling for social group and technical variables, as described above.

For each gene, we estimated the effect of timepoint and agonisms received on gene expression levels across phases using the following model:

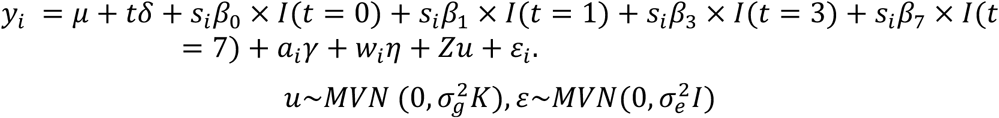

where *i* indexes the sample, and *y* is the *n* x 1 vector of residual gene expression levels for the *n* samples collected across four timepoints (0, 1, 3, and 7 days); μ is the intercept; t is an n x 1 vector of timepoints (Day0, Day1, Day3, Day7) and δ is the effect size for each day, respectively; *I* is an indicator variable for the timepoint (Day0 = 0, Day1 = 1, Day3 = 3, Day7 = 7); *s* is an *n* x 1 vector of the rate of agonisms received, and β_0_, β_1_, β_3_ and β_7_ are their effect sizes; *a* is an *n* x 1 vector of age in years at Day0 and γ is its effect size; and *w* is an *n* x 1 vector of individual body mass and η is its effect size. The vector *u* is an *m* x 1 vector of random effects, with a covariance structure determined by the genetic relatedness among the *m* unique individuals in the sample (described by *K*, a matrix of pairwise relatedness estimates derived from genotypes at 62,648 SNVs) and the additive genetic variance in *y*. *Z* is an *n* x *m* matrix of 1’s and 0’s that maps gene expression measurements to unique individuals in *u* to control for the repeated sampling of the same animals across timepoints. Residual errors are represented by the *n* x 1 vector ε representing the environmental variance component, *I* is the identity matrix, and *MVN* denotes the multivariate normal distribution.

For each gene, we tested the null hypotheses that β_j_ = 0 for j = 0, 1, 3, 7 (i.e., that there is no effect of social behavior on gene expression) and δ = 0 (i.e., that there is no effect of timepoint on gene expression) versus the alternative hypotheses β_j_ ≠ 0 for j = 0, 1, 3, 7 and δ ≠ 0. We created empirical null distributions for social behavioral metrics by permuting the metric across individuals (blocked across all three timepoints) 1,000 times and rerunning the analysis for each of the permutations. Similarly, the timepoint empirical null was generated by randomizing the timepoint label within the data generated from each individual macaque 1,000 times. We calculated permutation-based FDRs for social behavioral metrics and treatment timepoint following a generalization of the false discovery rate method of Storey and Tibshirani using the empirical null p-value distributions (*137*).

### Antibody prediction

To identify genes that predict the antibody response to vaccination, we tested for associations between fold change in gene expression following vaccination (differential gene expression, DGE, at days 1, 3 and 7) and fold change in influenza vaccine-specific IgG titers detected 28 days after vaccination compared to pre-vaccination baseline (FC Ab d28). Differential gene expression at a given timepoint was calculated as the difference between the residuals from batch-corrected expression values at days 1, 3 or 7 after vaccination and those at baseline (day 0). For each gene and each timepoint at days 1, 3, and 7, we fitted the following linear regression model, controlling for age and weight:

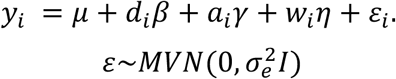

where *i* indexes the study subject, and *y* is the *n* x 1 vector of log-transformed fold changes in vaccine-specific IgG titers from day 28 to day 0, *d_i_* is the differential gene expression at a given timepoint, corrected for batch effects and β its effect size. The remaining notations follow the model for social rank effects on the antibody response, above. To correct for multiple testing, we produced empirical null distributions for the antibody response by permuting FC Ab d28 values among individuals 100 times. We used the resulting p-value distribution as an empirical null distribution to calculate false discovery rates using q-values ((*138*), R package qvalue version 2.4.2).

### Hierarchical clustering of vaccine-responsive genes

To identify shared trajectories of gene expression in the context of immune responses to influenza vaccination we performed unsupervised hierarchical clustering on the effect sizes (expressed in fold change) of genes that significantly responded to vaccination (1% FDR, n = 4,782). Standardized gene-level vaccine effects on days 1, 3 and 7 were first scaled so that each gene had sd = 1 across all samples, standardizing changes in expression, and then clustered based on R implementation of Ward’s (D2) minimum variance method. The optimal number of gene clusters was estimated based on within-cluster sum of squares using the *factoextra* R package (*139*)

### Enrichment analysis

We conducted gene set enrichment analysis (GSEA) using pre-ranked gene lists and over-representation analysis on subsets of significant genes to test for enrichment in biological functions. Specifically, we assessed enrichment in C5 gene ontology (GO) biological processes and the C2 Reactome subset of canonical pathways from the Molecular Signatures Database (MSigDB) (*140*). These gene sets were mapped from human to rhesus macaque orthologs and filtered to include only those present in our dataset. As input for GSEA, we ranked genes in the agonisms received-gene expression data set by standardized effect size from most negative (more highly expressed in animals who were rarely targeted by agonistic behavior and were typically high ranking) to most positive (more highly expressed in animals who were frequently targeted in agonisms, who were typically low ranking; Data S7). For genes in the gene expression-antibody titer analysis, we ranked genes by standardized effect size from most positive (cases in which large fold-changes in gene expression levels positively predict fold change in antibody titers) to more negative (cases in which large fold-changes in gene expression levels negatively predict fold change in antibody titers; Data S8). We also tested for over-representation of particular gene sets in subsets of vaccine-responsive genes that shared the same temporal expression dynamic (cluster genes), against a background of all expressed genes (Fig. 3A and Data S5). Analyses were carried out with clusterProfiler R package (*141*). P-values for the enrichment scores were adjusted for multiple comparisons with Benjamini-Hochberg (BH) correction.

Given the importance of the innate and adaptive immune responses in the response to vaccination, we were particularly interested in enrichment of genes involved in immune function. We therefore filtered the output of enrichment analyses for terms hierarchically related to broader immune categories in GO and Reactome collections of pathways. We used the GO.db R package (*142*) to select all descendant (offspring) GO terms deriving from “immune system process” (GO:0002376), “defense response” (GO:0006952), “cytokine production” (GO:0001816), “cytokine response” (GO:0034097), “response to virus” (GO:0009615) and “viral process” (GO:0016032). For Reactome pathways, the ReactomeContentService4R R package (*143*) was used to recursively retrieve participant pathways in “Innate Immune System” (R-HSA-168249), “Adaptive Immune System” (R-HSA-1280218), “Cytokine Signaling in Immune system” (R-HSA-1280215) and viral respiratory infections “Influenza Infection” (R-HSA-168255) and “SARS-CoV Infections” (R-HSA-9679506). Significant enrichments were reported as BH-corrected (adjusted) p-values at a 1% FDR threshold.

We calculated individual-level gene set pathway scores with the R package gsva (*101*), which computes sample-wise gene set enrichment scores as a function of genes inside and outside the gene set of interest. We generated pathway scores based on the expression of all genes associated with a given pathway that were present in our dataset. Pathway scores for immune pathways of interest were plotted against agonisms received (Fig. 3C) or fold increase in antibody titers (Fig. 4A).

## Acknowledgments

We thank J. Whitley, J. Bailey, and J. Johnson for maintaining the study subjects and collecting behavioral data; J. Guthmiller and the P. Wilson lab for guidance in implementing antibody assays and members of the L.B.B. and J.T. laboratories for helpful discussions. We thank the University of Chicago Genomics Facility (RRID: SCR_019196) for their assistance with sequencing. Figures 1A and 2A were created with BioRender.com.

## Funding

ENPRC base grant P51OD011132

National Institutes of Health grants R01GM102562 and R01AG057235 (VM, JT, LBB)

National Institutes of Health fellowship F32AG067704 (CRC)

National Institutes of Health fellowship F32AG062120 (NS)

National Institutes of Health fellowship F32AG064883 (PM)

## Author contributions

Conceptualization: VM, JT and LBB.

Methodology: JBB, CRC, NDS, PLM and RAG.

Investigation: JBB, CRC, AD, MS, TM and CB.

Visualization: JBB.

Supervision: VM, JT and LBB.

Writing—original draft: JBB, CRC, JT and LBB.

Writing—review & editing: JBB, CRC, JT and LBB.

## Competing interests

Authors declare that they have no competing interests.

## Data and materials availability

All data needed to evaluate the conclusions in the paper are present in the paper and/or the Supplementary Materials. The gene expression data reported in this paper has been deposited into the NCBI Gene Expression Omnibus (GEO) with accession number GSE318613. Relevant code and materials needed to reproduce the main results can be found on Zenodo.

## Supplementary Materials

**Fig. S1.**
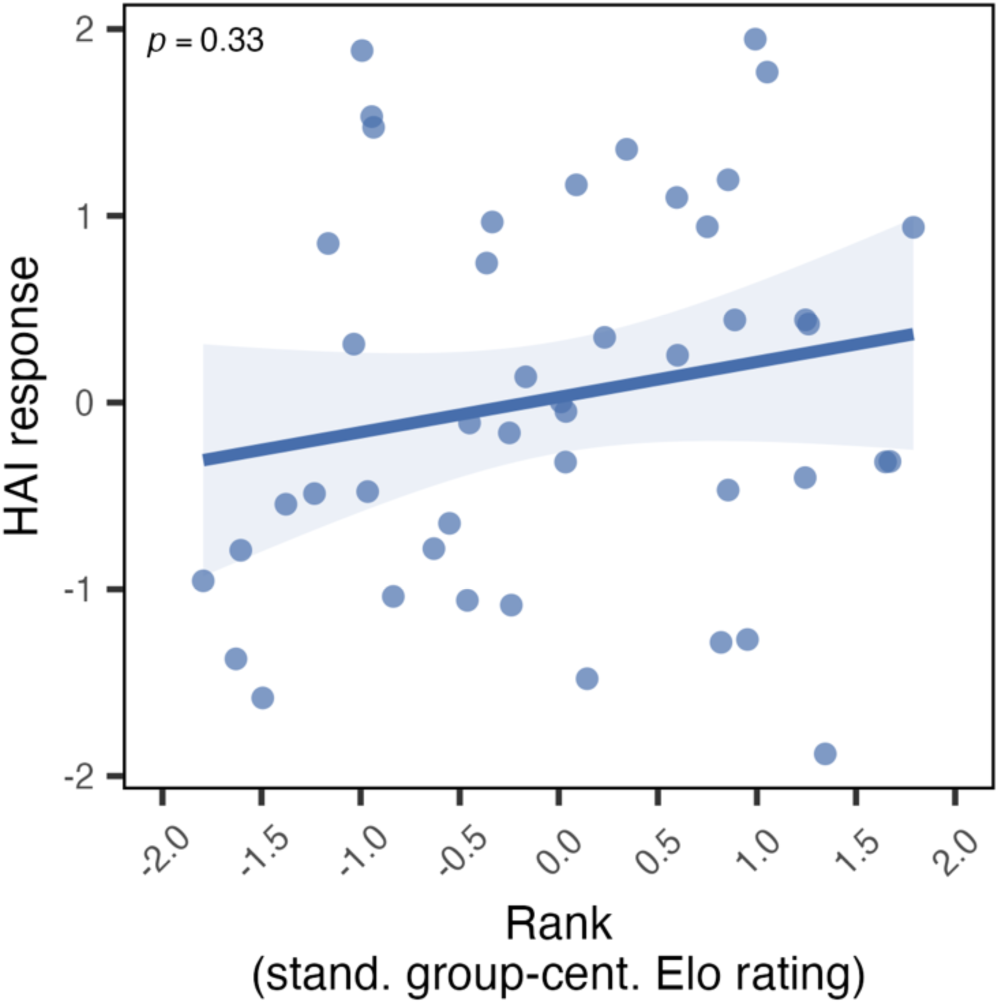
Effect of dominance rank, measured by Elo rating, on the HAI antibody response. (linear mixed model *p* = 0.33). Partial residuals plot for HAI titers (y-axis) as a function of dominance rank, adjusted for the contribution of age, body mass, group membership, and kinship.

**Fig. S2.**
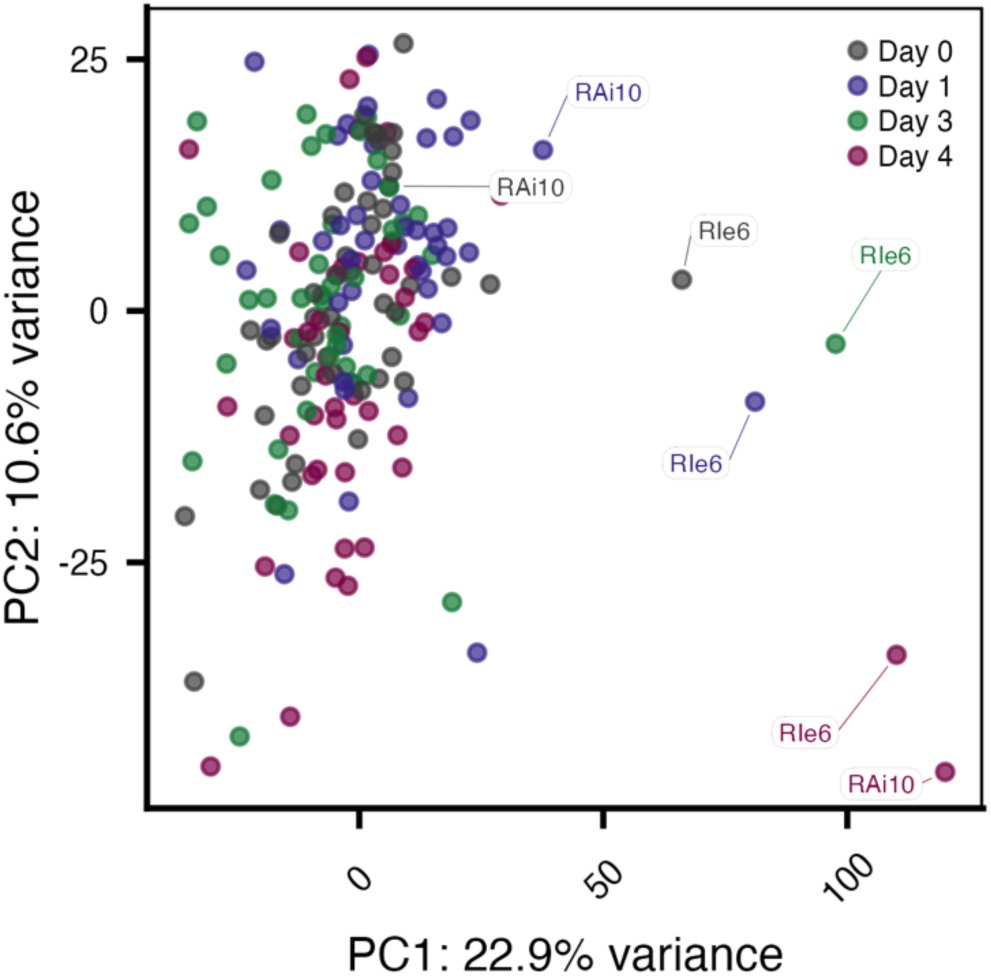
Principal component analysis of gene expression data across all 4 timepoints. We excluded two individuals who were conspicuous outliers (after batch correction) from subsequent analysis. Samples from the two individuals (Ai10 and Ie6) are labeled above.

**Fig. S3.**
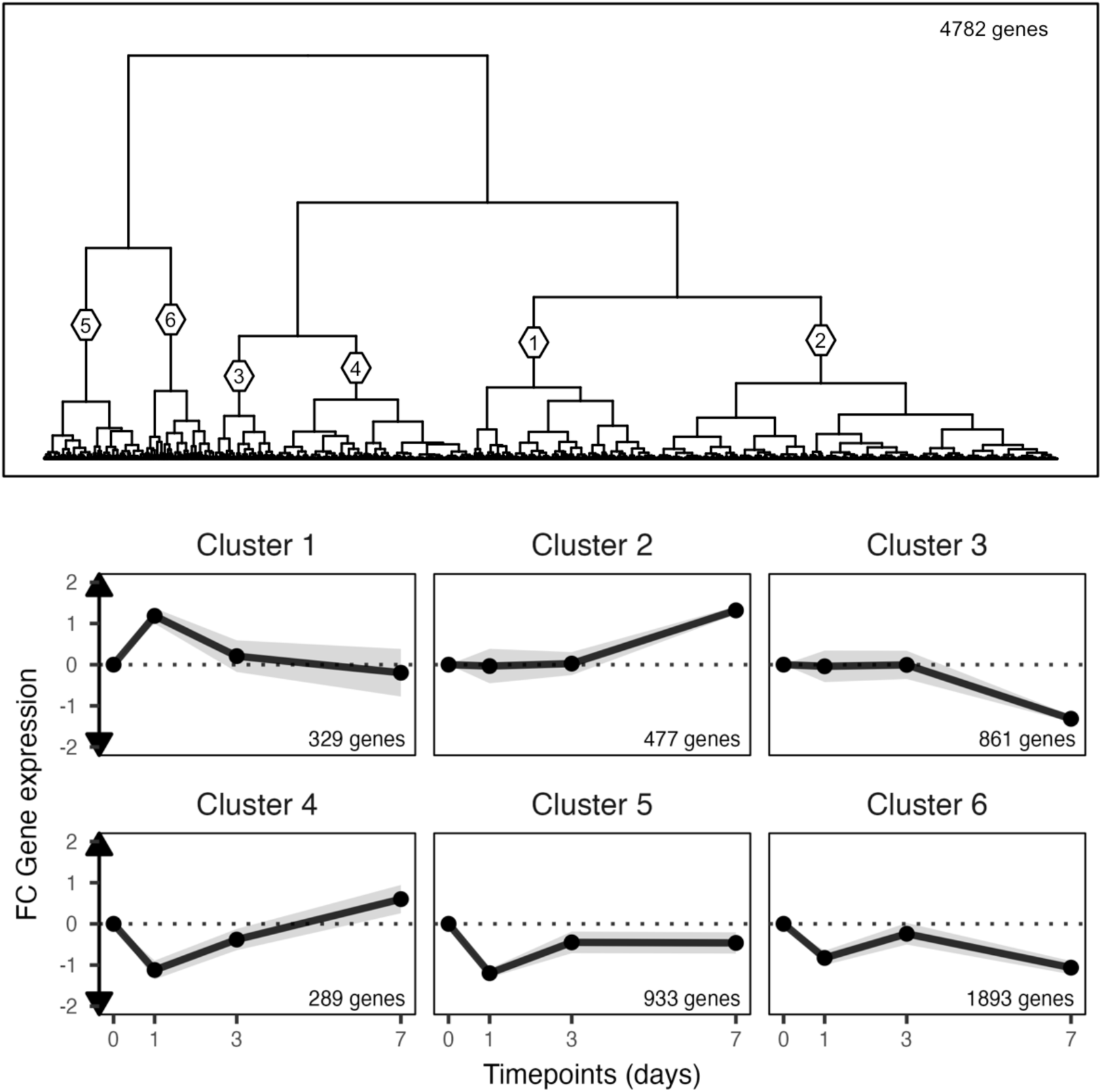
Hierarchical clustering of genes strongly responsive to the vaccine (FDR 1%). Unsupervised clustering of differentially expressed genes upon vaccination into 6 gene clusters that share similar transcriptional temporal dynamics. We focus on Clusters 1, 2, and 3 in the main text because we identified no enrichments for immune processes in Clusters 4, 5, or 6.

**Fig. S4.**
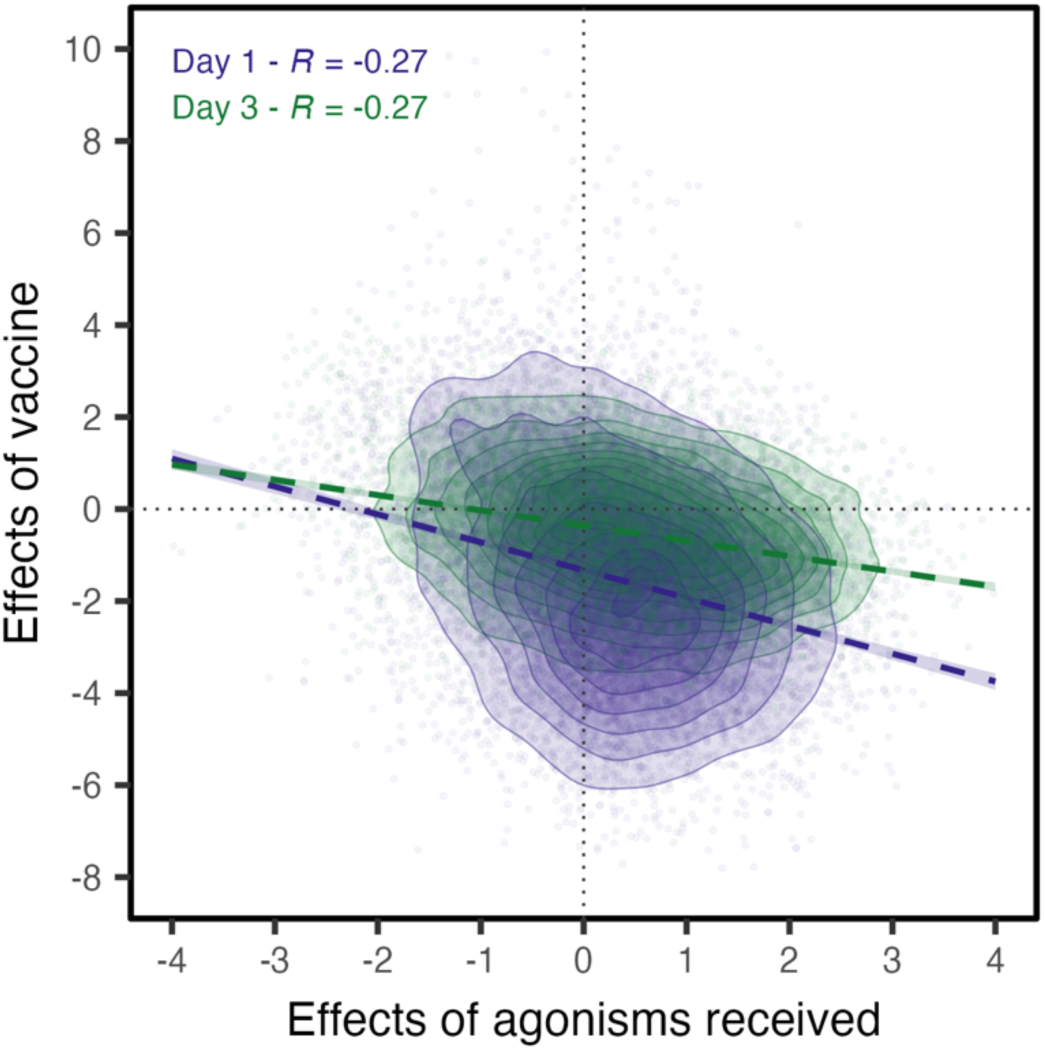
Correlation between the effects of influenza vaccine and agonisms received on gene expression. Correlation of gene-level effect sizes (standardized betas) between the vaccine response and agonisms received at days 1 and 3 following vaccination (day 7 shows the strongest correlation; see Figure 3B).

**Fig. S5.**
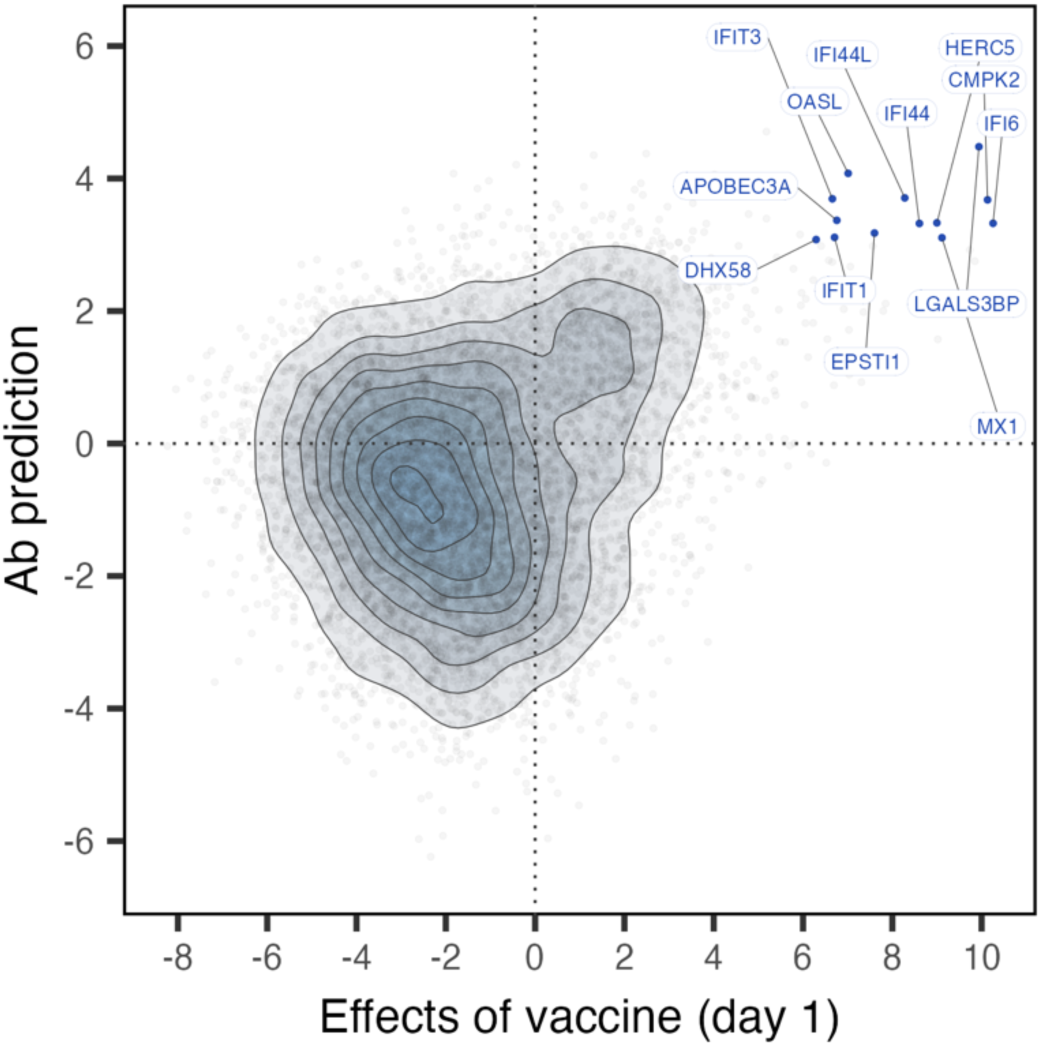
Correlation between effects of influenza vaccine and antibody prediction. Correlation of gene-level effect sizes (standardized betas) between the vaccine response on day 1 and the relationship between gene expression fold change and the fold change antibody response from baseline to day 28. ISG genes that positively predict the antibody response and that are strongly upregulated 1 day after vaccination tend to fall in the extreme upper right quadrant and are labeled with the gene name.

**Data S1 (Microsoft Excel Format).** Study subject information and behavioral data.

**Data S2 (Microsoft Excel Format).** Titers of vaccine-specific IgG antibodies (IgG, log_10_) and haemagglutination-protective serum antibodies (HAI, log_2_) for individuals in phases 1 (IgG, HAI) and 2 (IgG).

**Data S3 (Microsoft Excel Format).** Full parameter estimates, p-values, and FDR-corrected p-values by gene for flu vaccine effects per timepoint.

**Data S4 (Microsoft Excel Format).** Assignments of vaccine responsive genes (FDR 1%) to clusters that share similar temporal gene expression dynamics post-vaccination, as determined by hierarchical clustering.

**Data S5 (Microsoft Excel Format).** Over-representation analysis results (FDR 1%) - clusters of genes that respond to influenza vaccine (differentially expressed genes, FDR 1%).

**Data S6 (Microsoft Excel Format).** Full parameter estimates, p-values, and FDR-corrected p-values by gene for agonism received effects, within vaccine timepoint.

**Data S7 (Microsoft Excel Format).** Gene set enrichment analysis results (FDR 1%) for the effects of agonisms received at different timepoints post-vaccination.

**Data S8 (Microsoft Excel Format).** Gene set enrichment analysis results (FDR 1%) for genes where the fold-change response to vaccination predicts the foldchange antibody response at day 28 post-vaccination.

## Notes

https://www.ncbi.nlm.nih.gov/geo/query/acc.cgi?acc=GSE318613

### Competing Interest Statement

The authors have declared no competing interest.

## References

1. M. Marmot, Social determinants of health inequalities. Lancet Lond. Engl. 365, 1099–1104 (2005).

2. J. Holt-Lunstad, T. B. Smith, J. B. Layton, Social relationships and mortality risk: a meta-analytic review. PLoS Med. 7, e1000316 (2010).

3. J. Holt-Lunstad, T. B. Smith, M. Baker, T. Harris, D. Stephenson, Loneliness and social isolation as risk factors for mortality: a meta-analytic review. Perspect. Psychol. Sci. J. Assoc. Psychol. Sci. 10, 227–237 (2015).

4. S. Stringhini, C. Carmeli, M. Jokela, M. Avendaño, P. Muennig, F. Guida, F. Ricceri, A. d’Errico, H. Barros, M. Bochud, M. Chadeau-Hyam, F. Clavel-Chapelon, G. Costa, C. Delpierre, S. Fraga, M. Goldberg, G. G. Giles, V. Krogh, M. Kelly-Irving, R. Layte, A. M. Lasserre, M. G. Marmot, M. Preisig, M. J. Shipley, P. Vollenweider, M. Zins, I. Kawachi, A. Steptoe, J. P. Mackenbach, P. Vineis, M. Kivimäki, LIFEPATH consortium, Socioeconomic status and the 25 × 25 risk factors as determinants of premature mortality: a multicohort study and meta-analysis of 1·7 million men and women. Lancet Lond. Engl. 389, 1229–1237 (2017).

5. V. J. Felitti, R. F. Anda, D. Nordenberg, D. F. Williamson, A. M. Spitz, V. Edwards, M. P. Koss, J. S. Marks, Relationship of childhood abuse and household dysfunction to many of the leading causes of death in adults. The Adverse Childhood Experiences (ACE) Study. Am. J. Prev. Med. 14, 245–258 (1998).

6. P. Braveman, L. Gottlieb, The Social Determinants of Health: It’s Time to Consider the Causes of the Causes. Public Health Rep. 129, 19–31 (2014).

7. K. Steenland, S. Hu, J. Walker, All-cause and cause-specific mortality by socioeconomic status among employed persons in 27 US states, 1984-1997. Am. J. Public Health 94, 1037–1042 (2004).

8. A. Formanack, A. Doshi, R. Valdez, I. Williams, J. R. Moorman, P. Chernyavskiy, Race, Class, and Place Modify Mortality Rates for the Leading Causes of Death in the United States, 1999-2021. J. Gen. Intern. Med., 1–9 (2023).

9. W. M. Schultz, H. M. Kelli, J. C. Lisko, T. Varghese, J. Shen, P. Sandesara, A. A. Quyyumi, H. A. Taylor, M. Gulati, J. G. Harold, J. H. Mieres, K. C. Ferdinand, G. A. Mensah, L. S. Sperling, Socioeconomic Status and Cardiovascular Outcomes: Challenges and Interventions. Circulation 137, 2166–2178 (2018).

10. J. M. O’Connor, T. Sedghi, M. Dhodapkar, M. J. Kane, C. P. Gross, Factors Associated With Cancer Disparities Among Low-, Medium-, and High-Income US Counties. JAMA Netw. Open 1, e183146 (2018).

11. S. Saydah, K. Lochner, Socioeconomic status and risk of diabetes-related mortality in the U.S. Public Health Rep. Wash. DC 1974 125, 377–388 (2010).

12. N. E. Adler, J. M. Ostrove, Socioeconomic status and health: what we know and what we don’t. Ann. N. Y. Acad. Sci. 896, 3–15 (1999).

13. N. E. Adler, T. Boyce, M. A. Chesney, S. Cohen, S. Folkman, R. L. Kahn, S. L. Syme, Socioeconomic status and health. The challenge of the gradient. Am. Psychol. 49, 15–24 (1994).

14. K. A. Matthews, L. C. Gallo, Psychological perspectives on pathways linking socioeconomic status and physical health. Annu. Rev. Psychol. 62, 501–530 (2011).

15. S. Ravi, M. J. Shanahan, B. Levitt, K. M. Harris, S. W. Cole, Socioeconomic inequalities in early adulthood disrupt the immune transcriptomic landscape via upstream regulators. Sci. Rep. 14, 1–11 (2024).

16. S. E. Taylor, Mechanisms linking early life stress to adult health outcomes. Proc. Natl. Acad. Sci. 107, 8507–8512 (2010).

17. R. Glaser, B. Rabin, M. Chesney, S. Cohen, B. Natelson, Stress-induced immunomodulation: implications for infectious diseases? JAMA 281, 2268–2270 (1999).

18. G. Miller, E. Chen, S. W. Cole, Health psychology: developing biologically plausible models linking the social world and physical health. Annu. Rev. Psychol. 60, 501–524 (2009).

19. K. A. Muscatell, S. N. Brosso, K. L. Humphreys, Socioeconomic status and inflammation: a meta-analysis. Mol. Psychiatry 25, 2189–2199 (2020).

20. E. Berger, R. Castagné, M. Chadeau-Hyam, M. Bochud, A. d’Errico, M. Gandini, M. Karimi, M. Kivimäki, V. Krogh, M. Marmot, S. Panico, M. Preisig, F. Ricceri, C. Sacerdote, A. Steptoe, S. Stringhini, R. Tumino, P. Vineis, C. Delpierre, M. Kelly-Irving, Multi-cohort study identifies social determinants of systemic inflammation over the life course. Nat. Commun. 10, 773 (2019).

21. D. Baumeister, R. Akhtar, S. Ciufolini, C. M. Pariante, V. Mondelli, Childhood trauma and adulthood inflammation: a meta-analysis of peripheral C-reactive protein, interleukin-6 and tumour necrosis factor-α. Mol. Psychiatry 21, 642–649 (2016).

22. E. T. Klopack, E. M. Crimmins, S. W. Cole, T. E. Seeman, J. E. Carroll, Social stressors associated with age-related T lymphocyte percentages in older US adults: Evidence from the US Health and Retirement Study. Proc. Natl. Acad. Sci. U. S. A. 119, e2202780119 (2022).

23. G. E. Miller, E. Chen, A. K. Fok, H. Walker, A. Lim, E. F. Nicholls, S. Cole, M. S. Kobor, Low early-life social class leaves a biological residue manifested by decreased glucocorticoid and increased proinflammatory signaling. Proc. Natl. Acad. Sci. 106, 14716–14721 (2009).

24. D. E. Alley, T. E. Seeman, J. Ki Kim, A. Karlamangla, P. Hu, E. M. Crimmins, Socioeconomic status and C-reactive protein levels in the US population: NHANES IV. Brain. Behav. Immun. 20, 498–504 (2006).

25. E. B. Loucks, L. Pilote, J. W. Lynch, H. Richard, N. D. Almeida, E. J. Benjamin, J. M. Murabito, Life course socioeconomic position is associated with inflammatory markers: the Framingham Offspring Study. Soc. Sci. Med. 1982 71, 187–195 (2010).

26. T. L. Gruenewald, S. Cohen, K. A. Matthews, R. Tracy, T. E. Seeman, Association of socioeconomic status with inflammation markers in black and white men and women in the Coronary Artery Risk Development in Young Adults (CARDIA) study. Soc. Sci. Med. 69, 451–459 (2009).

27. S. Stringhini, G. D. Batty, P. Bovet, M. J. Shipley, M. G. Marmot, M. Kumari, A. G. Tabak, M. Kivimäki, Association of lifecourse socioeconomic status with chronic inflammation and type 2 diabetes risk: the Whitehall II prospective cohort study. PLoS Med. 10, e1001479 (2013).

28. D. Gimeno, E. J. Brunner, G. D. O. Lowe, A. Rumley, M. G. Marmot, J. E. Ferrie, Adult socioeconomic position, C-reactive protein and interleukin-6 in the Whitehall II prospective study. Eur. J. Epidemiol. 22, 675–683 (2007).

29. M. G. Marmot, G. D. Smith, S. Stansfeld, C. Patel, F. North, J. Head, I. White, E. Brunner, A. Feeney, Health inequalities among British civil servants: the Whitehall II study. Lancet Lond. Engl. 337, 1387–1393 (1991).

30. A. Rosengren, A. Smyth, S. Rangarajan, C. Ramasundarahettige, S. I. Bangdiwala, K. F. AlHabib, A. Avezum, K. Bengtsson Boström, J. Chifamba, S. Gulec, R. Gupta, E. U. Igumbor, R. Iqbal, N. Ismail, P. Joseph, M. Kaur, R. Khatib, I. M. Kruger, P. Lamelas, F. Lanas, S. A. Lear, W. Li, C. Wang, D. Quiang, Y. Wang, P. Lopez-Jaramillo, N. Mohammadifard, V. Mohan, P. K. Mony, P. Poirier, S. Srilatha, A. Szuba, K. Teo, A. Wielgosz, K. E. Yeates, K. Yusoff, R. Yusuf, A. H. Yusufali, M. W. Attaei, M. McKee, S. Yusuf, Socioeconomic status and risk of cardiovascular disease in 20 low-income, middle-income, and high-income countries: the Prospective Urban Rural Epidemiologic (PURE) study. Lancet Glob. Health 7, e748–e760 (2019).

31. A. Steptoe, P. Zaninotto, Lower socioeconomic status and the acceleration of aging: An outcome-wide analysis. Proc. Natl. Acad. Sci. U. S. A. 117, 14911–14917 (2020).

32. M. Marmot, Health equity in England: the Marmot review 10 years on. BMJ 368, m693 (2020).

33. Q. An, J. Prejean, K. McDavid Harrison, X. Fang, Association between community socioeconomic position and HIV diagnosis rate among adults and adolescents in the United States, 2005 to 2009. Am. J. Public Health 103, 120–126 (2013).

34. J. L. Hadler, K. Yousey-Hindes, A. Pérez, E. J. Anderson, M. Bargsten, S. R. Bohm, M. Hill, B. Hogan, M. Laidler, M. L. Lindegren, K. L. Lung, E. Mermel, L. Miller, C. Morin, E. Parker, S. M. Zansky, S. S. Chaves, Influenza-Related Hospitalizations and Poverty Levels - United States, 2010-2012. MMWR Morb. Mortal. Wkly. Rep. 65, 101–105 (2016).

35. M. Karmakar, P. M. Lantz, R. Tipirneni, Association of Social and Demographic Factors With COVID-19 Incidence and Death Rates in the US. JAMA Netw. Open 4, e2036462 (2021).

36. J. B. Dowd, A. E. Aiello, D. E. Alley, Socioeconomic disparities in the seroprevalence of cytomegalovirus infection in the US population: NHANES III. Epidemiol. Infect. 137, 58–65 (2009).

37. H. A. Beydoun, J. Dail, B. Ugwu, A. Boueiz, M. A. Beydoun, Socio-demographic and behavioral correlates of herpes simplex virus type 1 and 2 infections and co-infections among adults in the USA. Int. J. Infect. Dis. IJID Off. Publ. Int. Soc. Infect. Dis. 14S3, e154 (2010).

38. H. H. Balfour, F. Sifakis, J. A. Sliman, J. A. Knight, D. O. Schmeling, W. Thomas, Age-specific prevalence of Epstein-Barr virus infection among individuals aged 6-19 years in the United States and factors affecting its acquisition. J. Infect. Dis. 208, 1286–1293 (2013).

39. H. C. S. Meier, M. N. Haan, C. F. Mendes de Leon, A. M. Simanek, J. B. Dowd, A. E. Aiello, Early life socioeconomic position and immune response to persistent infections among elderly Latinos. Soc. Sci. Med. 1982 166, 77–85 (2016).

40. A. Zajacova, J. B. Dowd, A. E. Aiello, Socioeconomic and race/ethnic patterns in persistent infection burden among U.S. adults. J. Gerontol. A. Biol. Sci. Med. Sci. 64, 272–279 (2009).

41. A. Pini, M. Stenbeck, I. Galanis, H. Kallberg, K. Danis, A. Tegnell, A. Wallensten, Socioeconomic disparities associated with 29 common infectious diseases in Sweden, 2005-14: an individually matched case-control study. Lancet Infect. Dis. 19, 165–176 (2019).

42. R. C. Stebbins, G. A. Noppert, A. E. Aiello, E. Cordoba, J. B. Ward, L. Feinstein, Persistent socioeconomic and racial and ethnic disparities in pathogen burden in the United States, 1999-2014. Epidemiol. Infect. 147, e301 (2019).

43. X. Ye, Y. Wang, Y. Zou, J. Tu, W. Tang, R. Yu, S. Yang, P. Huang, Associations of socioeconomic status with infectious diseases mediated by lifestyle, environmental pollution and chronic comorbidities: a comprehensive evaluation based on UK Biobank. Infect. Dis. Poverty 12, 5 (2023).

44. G. W. Evans, E. Kantrowitz, Socioeconomic status and health: the potential role of environmental risk exposure. Annu. Rev. Public Health 23, 303–331 (2002).

45. N. E. Adler, Judith Stewart, Health disparities across the lifespan: meaning, methods, and mechanisms. Ann. N. Y. Acad. Sci. 1186, 5–23 (2010).

46. A. Glatman-Freedman, K. Nichols, The effect of social determinants on immunization programs. Hum. Vaccines Immunother. 8, 293–301 (2012).

47. E. Cordoba, A. E. Aiello, Social Determinants of Influenza Illness and Outbreaks in the United States. N. C. Med. J. 77, 341–345 (2016).

48. V. Barry, S. Dasgupta, D. L. Weller, J. L. Kriss, B. L. Cadwell, C. Rose, C. Pingali, T. Musial, J. D. Sharpe, S. A. Flores, K. J. Greenlund, A. Patel, A. Stewart, J. R. Qualters, L. Harris, K. E. Barbour, C. L. Black, Patterns in COVID-19 Vaccination Coverage, by Social Vulnerability and Urbanicity - United States, December 14, 2020-May 1, 2021. MMWR Morb. Mortal. Wkly. Rep. 70, 818–824 (2021).

49. J. K. Kiecolt-Glaser, R. Glaser, S. Gravenstein, W. B. Malarkey, J. Sheridan, Chronic stress alters the immune response to influenza virus vaccine in older adults. Proc. Natl. Acad. Sci. U. S. A. 93, 3043–3047 (1996).

50. R. Glaser, J. K. Kiecolt-Glaser, W. B. Malarkey, J. F. Sheridan, The influence of psychological stress on the immune response to vaccines. Ann. N. Y. Acad. Sci. 840, 649–655 (1998).

51. K. Vedhara, N. K. Cox, G. K. Wilcock, P. Perks, M. Hunt, S. Anderson, S. L. Lightman, N. M. Shanks, Chronic stress in elderly carers of dementia patients and antibody response to influenza vaccination. The Lancet 353, 627–631 (1999).

52. R. Glaser, J. Sheridan, W. B. Malarkey, R. C. MacCallum, J. K. Kiecolt-Glaser, Chronic stress modulates the immune response to a pneumococcal pneumonia vaccine. Psychosom. Med. 62, 804–807 (2000).

53. S. Gallagher, A. C. Phillips, M. T. Drayson, D. Carroll, Caregiving for children with developmental disabilities is associated with a poor antibody response to influenza vaccination. Psychosom. Med. 71, 341–344 (2009).

54. S. Gallagher, A. C. Phillips, M. T. Drayson, D. Carroll, Parental caregivers of children with developmental disabilities mount a poor antibody response to pneumococcal vaccination. Brain. Behav. Immun. 23, 338–346 (2009).

55. A. A. Madison, M. R. Shrout, M. E. Renna, J. K. Kiecolt-Glaser, Psychological and Behavioral Predictors of Vaccine Efficacy: Considerations for COVID-19. Perspect. Psychol. Sci. 16, 191–203 (2021).

56. S. Cohen, Keynote presentation at the eight international congress of behavioral medicine Mainz, Germany August 25–28, 2004. Int. J. Behav. Med. 12, 123–131 (2005).

57. S. Cohen, D. A. Tyrrell, A. P. Smith, Psychological stress and susceptibility to the common cold. N. Engl. J. Med. 325, 606–612 (1991).

58. S. Cohen, S. Line, S. B. Manuck, B. S. Rabin, E. R. Heise, J. R. Kaplan, Chronic social stress, social status, and susceptibility to upper respiratory infections in nonhuman primates. Psychosom. Med. 59, 213–221 (1997).

59. S. Cohen, E. Frank, W. J. Doyle, D. P. Skoner, B. S. Rabin, J. M. Gwaltney, Types of stressors that increase susceptibility to the common cold in healthy adults. Health Psychol. Off. J. Div. Health Psychol. Am. Psychol. Assoc. 17, 214–223 (1998).

60. S. Cohen, Social status and susceptibility to respiratory infections. Ann. N. Y. Acad. Sci. 896, 246–253 (1999).

61. S. Cohen, W. J. Doyle, R. Turner, C. M. Alper, D. P. Skoner, Sociability and susceptibility to the common cold. Psychol. Sci. 14, 389–395 (2003).

62. S. Cohen, W. J. Doyle, R. B. Turner, C. M. Alper, D. P. Skoner, Childhood socioeconomic status and host resistance to infectious illness in adulthood. Psychosom. Med. 66, 553–558 (2004).

63. M. Bonaccio, A. Di Castelnuovo, G. Pounis, A. De Curtis, S. Costanzo, M. Persichillo, C. Cerletti, M. B. Donati, G. de Gaetano, L. Iacoviello, Moli-sani Study Investigators, Relative contribution of health-related behaviours and chronic diseases to the socioeconomic patterning of low-grade inflammation. Int. J. Public Health 62, 551–562 (2017).

64. M. Marmot, S. Friel, R. Bell, T. A. Houweling, S. Taylor, Closing the gap in a generation: health equity through action on the social determinants of health. The Lancet 372, 1661–1669 (2008).

65. R. R. Johnson, R. Storts, T. H. Welsh, C. J. R. Welsh, M. W. Meagher, Social stress alters the severity of acute Theiler’s virus infection. J. Neuroimmunol. 148, 74–85 (2004).

66. J. W. Mays, N. D. Powell, J. T. Hunzeker, M. L. Hanke, M. T. Bailey, J. F. Sheridan, Stress and the anti-influenza immune response: Repeated social defeat augments clonal expansion of CD8+T cells during primary influenza A viral infection. J. Neuroimmunol. 243, 34–42 (2012).

67. J. F. Sheridan, J. L. Stark, R. Avitsur, D. A. Padgett, Social disruption, immunity, and susceptibility to viral infection. Role of glucocorticoid insensitivity and NGF. Ann. N. Y. Acad. Sci. 917, 894–905 (2000).

68. A. Sommershof, M. Basler, C. Riether, H. Engler, M. Groettrup, Attenuation of the cytotoxic T lymphocyte response to lymphocytic choriomeningitis virus in mice subjected to chronic social stress. Brain. Behav. Immun. 25, 340–348 (2011).

69. D. L. Stark, J. W. Cauceglia, V. N. Sitzman, M. C. Repetto, J. M. Tadje, W. K. Potts, Friend virus severity is associated with male mouse social status and environmental temperature. Anim. Behav. 187, 221–231 (2022).

70. J. E. Cunnick, S. Cohen, B. S. Rabin, A. B. Carpenter, S. B. Manuck, J. R. Kaplan, Alterations in specific antibody production due to rank and social instability. Brain. Behav. Immun. 5, 357–369 (1991).

71. S. Cohen, J. R. Kaplan, J. E. Cunnick, S. B. Manuck, B. S. Rabin, Chronic Social Stress, Affiliation, and Cellular Immune Response in Nonhuman Primates. Psychol. Sci. 3, 301–305 (1992).

72. J. P. Capitanio, S. P. Mendoza, N. W. Lerche, W. A. Mason, Social stress results in altered glucocorticoid regulation and shorter survival in simian acquired immune deficiency syndrome. Proc. Natl. Acad. Sci. 95, 4714–4719 (1998).

73. E. K. Sloan, J. P. Capitanio, R. P. Tarara, S. P. Mendoza, W. A. Mason, S. W. Cole, Social stress enhances sympathetic innervation of primate lymph nodes: mechanisms and implications for viral pathogenesis. J. Neurosci. Off. J. Soc. Neurosci. 27, 8857–8865 (2007).

74. E. L. Kinnally, S. J. Martinez, K. Chun, J. P. Capitanio, L. C. Ceniceros, Early Social Stress Promotes Inflammation and Disease Risk in Rhesus Monkeys. Sci. Rep. 9, 7609 (2019).

75. S. M. Guerrero-Martin, L. H. Rubin, K. M. McGee, E. N. Shirk, S. E. Queen, M. Li, B. Bullock, B. W. Carlson, R. J. Adams, L. Gama, D. R. Graham, C. Zink, J. E. Clements, J. L. Mankowski, K. A. Metcalf Pate, Psychosocial Stress Alters the Immune Response and Results in Higher Viral Load During Acute Simian Immunodeficiency Virus Infection in a Pigtailed Macaque Model of Human Immunodeficiency Virus. J. Infect. Dis. 224, 2113–2121 (2021).

76. M. A. Pavez-Fox, J. E. Negron-Del Valle, I. J. Thompson, C. S. Walker, S. E. Bauman, O. Gonzalez, N. Compo, A. Ruiz-Lambides, M. I. Martinez, M. L. Platt, M. J. Montague, J. P. Higham, N. Snyder-Mackler, L. J. N. Brent, Sociality predicts individual variation in the immunity of free-ranging rhesus macaques. Physiol. Behav. 241, 113560 (2021).

77. A. Koert, A. Ploeger, C. L. H. Bockting, M. V. Schmidt, P. J. Lucassen, A. Schrantee, J. D. Mul, The social instability stress paradigm in rat and mouse: A systematic review of protocols, limitations, and recommendations. Neurobiol. Stress 15, 100410 (2021).

78. H. Jarrell, J. B. Hoffman, J. R. Kaplan, S. Berga, B. Kinkead, M. E. Wilson, Polymorphisms in the serotonin reuptake transporter gene modify the consequences of social status on metabolic health in female rhesus monkeys. Physiol. Behav. 93, 807–819 (2008).

79. J. Tung, L. B. Barreiro, Z. P. Johnson, K. D. Hansen, V. Michopoulos, D. Toufexis, K. Michelini, M. E. Wilson, Y. Gilad, Social environment is associated with gene regulatory variation in the rhesus macaque immune system. Proc. Natl. Acad. Sci. U. S. A. 109, 6490–6495 (2012).

80. N. Snyder-Mackler, J. Sanz, J. N. Kohn, J. F. Brinkworth, S. Morrow, A. O. Shaver, J.-C. Grenier, R. Pique-Regi, Z. P. Johnson, M. E. Wilson, L. B. Barreiro, J. Tung, Social status alters immune regulation and response to infection in macaques. Science 354, 1041–1045 (2016).

81. N. Snyder-Mackler, J. N. Kohn, L. B. Barreiro, Z. P. Johnson, M. E. Wilson, J. Tung, Social status drives social relationships in groups of unrelated female rhesus macaques. Anim. Behav. 111, 307–317 (2016).

82. V. Michopoulos, M. Higgins, D. Toufexis, M. E. Wilson, Social subordination produces distinct stress-related phenotypes in female rhesus monkeys. Psychoneuroendocrinology 37, 1071–1085 (2012).

83. J. N. Kohn, N. Snyder-Mackler, L. B. Barreiro, Z. P. Johnson, J. Tung, M. E. Wilson, Dominance rank causally affects personality and glucocorticoid regulation in female rhesus macaques. Psychoneuroendocrinology 74, 179–188 (2016).

84. V. Michopoulos, K. M. Reding, M. E. Wilson, D. Toufexis, Social subordination impairs hypothalamic-pituitary-adrenal function in female rhesus monkeys. Horm. Behav. 62, 389–399 (2012).

85. N. D. Simons, V. Michopoulos, M. Wilson, L. B. Barreiro, J. Tung, Agonism and grooming behaviour explain social status effects on physiology and gene regulation in rhesus macaques. Philos. Trans. R. Soc. B Biol. Sci. 377, 20210132 (2022).

86. A. J. Lea, M. Y. Akinyi, R. Nyakundi, P. Mareri, F. Nyundo, T. Kariuki, S. C. Alberts, E. A. Archie, J. Tung, Dominance rank-associated gene expression is widespread, sex-specific, and a precursor to high social status in wild male baboons. Proc. Natl. Acad. Sci. 115, E12163–E12171 (2018).

87. N. Snyder-Mackler, J. Sanz, J. N. Kohn, T. Voyles, R. Pique-Regi, M. E. Wilson, L. B. Barreiro, J. Tung, Social status alters chromatin accessibility and the gene regulatory response to glucocorticoid stimulation in rhesus macaques. Proc. Natl. Acad. Sci. 116, 1219–1228 (2019).

88. J. Sanz, P. L. Maurizio, N. Snyder-Mackler, N. D. Simons, T. Voyles, J. Kohn, V. Michopoulos, M. Wilson, J. Tung, L. B. Barreiro, Social history and exposure to pathogen signals modulate social status effects on gene regulation in rhesus macaques. Proc. Natl. Acad. Sci. 117, 23317–23322 (2020).

89. M. R. S. Rosado, N. Marzan-Rivera, M. M. Watowich, A. D. N.-D. Valle, P. Pantoja, M. A. Pavez-Fox, E. R. Siracusa, E. B. Cooper, J. E. N.-D. Valle, D. Phillips, A. Ruiz-Lambides, M. I. Martinez, M. J. Montague, M. L. Platt, J. P. Higham, L. J. N. Brent, C. A. Sariol, N. Snyder-Mackler, Immune cell composition varies by age, sex and exposure to social adversity in free-ranging Rhesus Macaques. GeroScience 46, 2107–2122 (2023).

90. K. Adams, K. Yousey-Hindes, C. H. Bozio, S. Jain, P. D. Kirley, I. Armistead, N. B. Alden, K. P. Openo, L. S. Witt, M. L. Monroe, S. Kim, A. Falkowski, R. Lynfield, M. McMahon, M. R. Hoffman, Y. P. Shaw, N. L. Spina, A. Rowe, C. B. Felsen, E. Licherdell, K. Lung, E. Shiltz, A. Thomas, H. K. Talbot, W. Schaffner, M. T. Crossland, K. P. Olsen, L. W. Chang, C. N. Cummings, M. W. Tenforde, S. Garg, J. L. Hadler, A. O’Halloran, Social Vulnerability, Intervention Utilization, and Outcomes in US Adults Hospitalized With Influenza. JAMA Netw. Open 7, e2448003 (2024).

91. R. Chandrasekhar, C. Sloan, E. Mitchel, D. Ndi, N. Alden, A. Thomas, N. M. Bennett, P. D. Kirley, M. Hill, E. J. Anderson, R. Lynfield, K. Yousey-Hindes, M. Bargsten, S. M. Zansky, K. Lung, M. Schroeder, M. Monroe, S. Eckel, T. M. Markus, C. N. Cummings, S. Garg, W. Schaffner, M. L. Lindegren, Social determinants of influenza hospitalization in the United States. Influenza Other Respir. Viruses 11, 479–488 (2017).

92. A. D’Adamo, A. Schnake-Mahl, P. H. Mullachery, M. Lazo, A. V. Diez Roux, U. Bilal, Health disparities in past influenza pandemics: A scoping review of the literature. SSM - Popul. Health 21, 101314 (2023).

93. K. H. Grantz, M. S. Rane, H. Salje, G. E. Glass, S. E. Schachterle, D. A. T. Cummings, Disparities in influenza mortality and transmission related to sociodemographic factors within Chicago in the pandemic of 1918. Proc. Natl. Acad. Sci. 113, 13839–13844 (2016).

94. M. A. Chen, A. S. LeRoy, M. Majd, J. Y. Chen, R. L. Brown, L. M. Christian, C. P. Fagundes, Immune and Epigenetic Pathways Linking Childhood Adversity and Health Across the Lifespan. Front. Psychol. 12 (2021).

95. S. W. Cole, The Conserved Transcriptional Response to Adversity. Curr. Opin. Behav. Sci. 28, 31–37 (2019).

96. N. D. Simons, J. Tung, Social Status and Gene Regulation: Conservation and Context Dependence in Primates. Trends Cogn. Sci. 23, 722–725 (2019).

97. C. Neumann, J. Duboscq, C. Dubuc, A. M. Ginting, A. Engelhardt, Assessing dominance hierarchies: validation and advantages of progressive evaluation with Elo-rating. Anim. Behav. 82, 911–921 (2011).

98. S. W. Cole, Human social genomics. PLoS Genet. 10, e1004601 (2014).

99. H. I. Nakaya, T. Hagan, S. S. Duraisingham, E. K. Lee, M. Kwissa, N. Rouphael, D. Frasca, M. Gersten, A. K. Mehta, R. Gaujoux, G.-M. Li, S. Gupta, R. Ahmed, M. J. Mulligan, S. Shen-Orr, B. B. Blomberg, S. Subramaniam, B. Pulendran, Systems Analysis of Immunity to Influenza Vaccination across Multiple Years and in Diverse Populations Reveals Shared Molecular Signatures. Immunity 43, 1186–1198 (2015).

100. A. Lane, H. Q. Quach, I. G. Ovsyannikova, R. B. Kennedy, T. M. Ross, T. Einav, High-resolution antibody dynamics following influenza vaccination reveal predominantly weak responses as well as infrequent but durable immunity across the 2014-2022 seasons. Vaccine 64, 127677 (2025).

101. S. Hänzelmann, R. Castelo, J. Guinney, GSVA: gene set variation analysis for microarray and RNA-Seq data. BMC Bioinformatics 14, 7 (2013).

102. K. L. Bucasas, L. M. Franco, C. A. Shaw, M. S. Bray, J. M. Wells, D. Niño, N. Arden, J. M. Quarles, R. B. Couch, J. W. Belmont, Early Patterns of Gene Expression Correlate With the Humoral Immune Response to Influenza Vaccination in Humans. J. Infect. Dis. 203, 921–929 (2011).

103. H. I. Nakaya, J. Wrammert, E. K. Lee, L. Racioppi, S. Marie-Kunze, W. N. Haining, A. R. Means, S. P. Kasturi, N. Khan, G.-M. Li, M. McCausland, V. Kanchan, K. E. Kokko, S. Li, R. Elbein, A. K. Mehta, A. Aderem, K. Subbarao, R. Ahmed, B. Pulendran, Systems biology of vaccination for seasonal influenza in humans. Nat. Immunol. 12, 786–795 (2011).

104. L. M. Franco, K. L. Bucasas, J. M. Wells, D. Niño, X. Wang, G. E. Zapata, N. Arden, A. Renwick, P. Yu, J. M. Quarles, M. S. Bray, R. B. Couch, J. W. Belmont, C. A. Shaw, Integrative genomic analysis of the human immune response to influenza vaccination. eLife 2, e00299 (2013).

105. R. G. Cao, N. M. Suarez, G. Obermoser, S. M. C. Lopez, E. Flano, S. E. Mertz, R. A. Albrecht, A. García-Sastre, A. Mejias, H. Xu, H. Qin, D. Blankenship, K. Palucka, V. Pascual, O. Ramilo, Differences in Antibody Responses Between Trivalent Inactivated Influenza Vaccine and Live Attenuated Influenza Vaccine Correlate With the Kinetics and Magnitude of Interferon Signaling in Children. J. Infect. Dis. 210, 224–233 (2014).

106. L. M. Howard, K. L. Hoek, J. B. Goll, P. Samir, A. Galassie, T. M. Allos, X. Niu, L. E. Gordy, C. B. Creech, N. Prasad, T. L. Jensen, H. Hill, S. E. Levy, S. Joyce, A. J. Link, K. M. Edwards, Cell-Based Systems Biology Analysis of Human AS03-Adjuvanted H5N1 Avian Influenza Vaccine Responses: A Phase I Randomized Controlled Trial. PLoS ONE 12, e0167488 (2017).

107. P. Zimmermann, N. Curtis, Factors That Influence the Immune Response to Vaccination. Clin. Microbiol. Rev. 32, 10.1128/cmr.00084-18 (2019).

108. E. Tuaillon, A. Pisoni, N. Veyrenche, S. Rafasse, C. Niel, N. Gros, D. Muriaux, M.-C. Picot, S. Aouinti, P. Van de Perre, J. Bousquet, H. Blain, Antibody response after first and second BNT162b2 vaccination to predict the need for subsequent injections in nursing home residents. Sci. Rep. 12, 13749 (2022).

109. G. L. Salvagno, B. M. Henry, G. Lippi, The strength of association between pre-and post-booster BNT162b2 anti-SARS-CoV-2 antibodies levels depends on the immunoassay. Int. J. Infect. Dis. IJID Off. Publ. Int. Soc. Infect. Dis. 111, 65–67 (2021).

110. T. Perkmann, N. Perkmann-Nagele, P. Mucher, A. Radakovics, M. Repl, T. Koller, G. Jordakieva, O. F. Wagner, C. J. Binder, H. Haslacher, Initial SARS-CoV-2 vaccination response can predict booster response for BNT162b2 but not for AZD1222. Int. J. Infect. Dis. IJID Off. Publ. Int. Soc. Infect. Dis. 110, 309–313 (2021).

111. J. Janeway, Charles A., P. Travers, M. Walport, M. J. Shlomchik, “Immunological memory” in Immunobiology: The Immune System in Health and Disease. 5th Edition (Garland Science, 2001; https://www-ncbi-nlm-nih-gov.proxy.uchicago.edu/books/NBK27158/).

112. M. B. Azad, Y. Lissitsyn, G. E. Miller, A. B. Becker, K. T. HayGlass, A. L. Kozyrskyj, Influence of Socioeconomic Status Trajectories on Innate Immune Responsiveness in Children. PLoS ONE 7, e38669 (2012).

113. A. Gugushvili, G. Bulczak, O. Zelinska, J. Koltai, Socioeconomic position, social mobility, and health selection effects on allostatic load in the United States. PLOS ONE 16, e0254414 (2021).

114. P. H. Lam, E. Chen, J. J. Chiang, G. E. Miller, Socioeconomic disadvantage, chronic stress, and proinflammatory phenotype: an integrative data analysis across the lifecourse. PNAS Nexus 1, pgac219 (2022).

115. N. Na-Ek, P. Demakakos, Social mobility and inflammatory and metabolic markers at older ages: the English Longitudinal Study of Ageing. J. Epidemiol. Community Health 71, 253–260 (2017).

116. Y. C. Yang, K. Gerken, K. Schorpp, C. Boen, K. M. Harris, Early-Life Socioeconomic Status and Adult Physiological Functioning: A Life Course Examination of Biosocial Mechanisms. Biodemography Soc. Biol. 63, 87–103 (2017).

117. F. Diaz-Mitoma, I. Alvarez-Maya, A. Dabrowski, J. Jaffey, R. Frost, S. Aucoin, M. Kryworuchko, M. Lapner, H. Tadesse, A. Giulivi, Transcriptional analysis of human peripheral blood mononuclear cells after influenza immunization. J. Clin. Virol. 31, 100–112 (2004).

118. E. A. Voigt, D. E. Grill, M. T. Zimmermann, W. L. Simon, I. G. Ovsyannikova, R. B. Kennedy, G. A. Poland, Transcriptomic signatures of cellular and humoral immune responses in older adults after seasonal influenza vaccination identified by data-driven clustering. Sci. Rep. 8, 739 (2018).

119. C. Thomas, R. Tampé, MHC I chaperone complexes shaping immunity. Curr. Opin. Immunol. 58, 9–15 (2019).

120. J. Wang, J. Lee, D. Liem, P. Ping, HSPA5 Gene Encoding Hsp70 Chaperone BiP in the Endoplasmic Reticulum. Gene 618, 14–23 (2017).

121. M. Zehner, A. L. Marschall, E. Bos, J.-G. Schloetel, C. Kreer, D. Fehrenschild, A. Limmer, F. Ossendorp, T. Lang, A. J. Koster, S. Dübel, S. Burgdorf, The Translocon Protein Sec61 Mediates Antigen Transport from Endosomes in the Cytosol for Cross-Presentation to CD8+ T Cells. Immunity 42, 850–863 (2015).

122. B. Alberts, A. Johnson, J. Lewis, M. Raff, K. Roberts, P. Walter, “Helper T Cells and Lymphocyte Activation” in Molecular Biology of the Cell. 4th Edition (Garland Science, 2002; https://www.ncbi.nlm.nih.gov/books/NBK26827/).

123. J. Hoes, A. G. C. Boef, M. J. Knol, H. E. de Melker, L. Mollema, F. R. M. van der Klis, N. Y. Rots, D. van Baarle, Socioeconomic Status Is Associated With Antibody Levels Against Vaccine Preventable Diseases in the Netherlands. Front. Public Health 6, 209 (2018).

124. F. S. Laroux, Mechanisms of inflammation: the good, the bad and the ugly. Front. Biosci. J. Virtual Libr. 9, 3156–3162 (2004).

125. D. S. Sade, “A Longitudinal Study of Social Behavior of Rhesus Monkeys” in The Functional and Evolutionary Biology of Primates (Routledge, 1972).

126. C. Neumann, L. Kulik, EloRating: Animal Dominance Hierarchies by Elo Rating, (2024); https://CRAN.R-project.org/package=EloRating.

127. E. Spackman, I. Sitaras, “Hemagglutination Inhibition Assay” in Animal Influenza Virus: Methods and Protocols, E. Spackman, Ed. (Springer US, New York, NY, 2020; 10.1007/978-1-0716-0346-8_2), pp. 11–28.

128. W. E. P. Beyer, A. M. Palache, G. Lüchters, J. Nauta, A. D. M. E. Osterhaus, Seroprotection rate, mean fold increase, seroconversion rate: which parameter adequately expresses seroresponse to influenza vaccination? Virus Res. 103, 125–132 (2004).

129. T. M. Therneau, Coxme: Mixed Effects Cox Models (2024; https://CRAN.R-project.org/package=coxme).

130. A. M. Bolger, M. Lohse, B. Usadel, Trimmomatic: a flexible trimmer for Illumina sequence data. Bioinformatics 30, 2114–2120 (2014).

131. A. Dobin, C. A. Davis, F. Schlesinger, J. Drenkow, C. Zaleski, S. Jha, P. Batut, M. Chaisson, T. R. Gingeras, STAR: ultrafast universal RNA-seq aligner. Bioinforma. Oxf. Engl. 29, 15–21 (2013).

132. G. H. Putri, S. Anders, P. T. Pyl, J. E. Pimanda, F. Zanini, Analysing high-throughput sequencing data in Python with HTSeq 2.0. Bioinformatics 38, 2943–2945 (2022).

133. R Core Team, R: A Language and Environment for Statistical Computing (R Foundation for Statistical Computing, Vienna, Austria, 2021; https://www.R-project.org/).

134. M. E. Ritchie, B. Phipson, D. Wu, Y. Hu, C. W. Law, W. Shi, G. K. Smyth, limma powers differential expression analyses for RNA-sequencing and microarray studies. Nucleic Acids Res. 43, e47 (2015).

135. M. Lipatov, K. Sanjeev, R. Patro, K. R. Veeramah, Maximum Likelihood Estimation of Biological Relatedness from Low Coverage Sequencing Data. bioRxiv [Preprint] (2015). 10.1101/023374.

136. D. Akdemir, O. U. Godfrey, EMMREML: Fitting Mixed Models with Known Covariance Structures, version 3.1 (2015); https://CRAN.R-project.org/package=EMMREML.

137. J. D. Storey, R. Tibshirani, Statistical significance for genomewide studies. Proc. Natl. Acad. Sci. U. S. A. 100, 9440–9445 (2003).

138. J. D. Storey, A. J. Bass, A. Dabney, D. Robinson, qvalue: Q-value estimation for false discovery rate control, (2024); 10.18129/B9.bioc.qvalue.

139. A. Kassambara, F. Mundt, factoextra: Extract and Visualize the Results of Multivariate Data Analyses, version 1.0.7 (2020); https://CRAN.R-project.org/package=factoextra.

140. A. Subramanian, P. Tamayo, V. K. Mootha, S. Mukherjee, B. L. Ebert, M. A. Gillette, A. Paulovich, S. L. Pomeroy, T. R. Golub, E. S. Lander, J. P. Mesirov, Gene set enrichment analysis: A knowledge-based approach for interpreting genome-wide expression profiles. Proc. Natl. Acad. Sci. 102, 15545–15550 (2005).

141. T. Wu, E. Hu, S. Xu, M. Chen, P. Guo, Z. Dai, T. Feng, L. Zhou, W. Tang, L. Zhan, X. Fu, S. Liu, X. Bo, G. Yu, clusterProfiler 4.0: A universal enrichment tool for interpreting omics data. Innov. Camb. Mass 2, 100141 (2021).

142. M. Carlson, GO.db: A set of annotation maps describing the entire Gene Ontology, version 3.19.1 (2019); https://bioconductor.org/packages/GO.db/.

143. C. L. Poon, J. Cook, S. Shorser, J. Weiser, R. Haw, L. Stein, R interface to the reactome graph database, volume 10, version 1.12.0 (2021); doi:10.7490/f1000research.1118690.1.

